# yakRNA Design: A semantic multimodal RNA composer

**DOI:** 10.64898/2026.04.22.720245

**Authors:** Louie N. Pinpin, Yousuf A. Khan

## Abstract

Like proteins, RNAs have been a target for exploitation to generate synthetic molecules that can adopt enhanced and novel functions^1^. High-throughput assays^2^, combined with meticulous biochemistry, have led to the generation of some artificial RNAs but the ability to algorithmically program RNAs with an intended function remains difficult. While generative models have revolutionized protein design^3,4^, RNA design remains challenging due to the dearth of 3D RNA structures. Here we present You Always Know RNA (yakRNA) Design, a frontier language model that can simultaneously reason over sequence, biological function, consensus sequence, and secondary structure to generate functional RNAs. yakRNA Design is not trained on any 3D structural information but rather co-learned over a corpus of semantically labeled sequences that we designed. yakRNA Design, without any human intervention, fine-tuning, sequence optimization, scoring or selection, zero-shot designed 17 (out of 84 total designs) synthetic RNAs that were able to efficiently induce ribosomes to change their reading frames during elongation at rates comparable to or higher than most natural sequences. One of these efficient synthetic RNAs was found to have no identity to any sequence in the known universe, demonstrating how a model with semantic understanding of RNA has rich generative capabilities.

---

The immense amount of global collaboration in solving protein structures led to an explosion of protein structures in the protein data bank (PDB)^5^, which allowed for the revolution in protein structure prediction with machine learning^6^ and now in protein design^3,4^; the conception of novel protein sequences with potentially new or enhanced biochemical function.

RNA design poses as an attractive target for advancements in therapeutics, drug designs, biosensors, and binders due to RNA’s inherent ease of programmability and synthesis. Indeed, specialized RNAs are already utilized as CRISPR single guide RNAs (sgRNAs)^7^, antisense oligonucleotides (ASOs)^8^, small interfering RNAs (siRNAs)^9^, scaffolded assemblies for intracellular reactions^10^, and riboswitches for metabolic engineering^11^.

However, despite the biochemical, biological, and clinical characterization of the importance of RNA structure, less than 1% of the PDB consist of RNA-only entries providing limiting training data^12,13^. This has complicated the ability to reliably predict RNA structure^14^ and hindered the development of RNA design. While some machine learning based structured RNA design tools exist^15–17^, they rely on the PDB as a source of training data which, given how sparse the amount of RNA structures that exist, leads to brittleness and models that lack generalizability^18^. Thus, to address the limited amount of training data for RNA, innovative approaches are demanded.

Others have trained RNA foundational models purely on raw corpuses of unannotated sequences^19–22^, which have showed promise in representational learning of RNA, are limited in their generative capabilities and typically require fine-tuning and/or additional training for specific tasks, limiting their adoption and useability. Furthermore, creating the largest database of unlabeled RNA sequences may not be the most effective strategy since increasing database size for RNAs does not necessarily correlate to improved downstream effectiveness at a task^23^.

Multimodal co-learning systems learn multiple different modalities at once, are typically associated with text, audio, video, etc and show better performance than typical unimodal (e.g. single modality) systems^24^.

Here we present yakRNA Design, an RNA composer. yakRNA Design was not trained on any 3D RNA structural information. Rather than being constrained by the limited of RNA structural data and following the roadmap of synthetic biology in proteins (accurate protein structure prediction begetting generative protein design), we sidestepped this traditional roadmap by instead devising a semantically labeled dataset in which each piece of training data contains an RNA sequence, a corresponding secondary structure, a consensus representation created from similar RNAs, and a prescribed biological function. This database, termed semanticRNAFamilies (semRfam), was used as the basis for training yakRNA Design so that it could co-learn the relationship between these different streams of information in a single unified vocabulary and gain a richer understanding of RNA, as opposed to unimodal models^25^.

yakRNA Design can take any combination of these modalities, based on the user’s preference, to generate synthetic RNAs that meet the functional specifications of the user. We also show that incorporating different prompting modalities together in a variety of combinations improves the ability of yakRNA Design to create sequences that align to the user’s preference, indicating synergy of the modalities for generating synthetic RNAs.

Lastly, yakRNA Design directly zero-shot synthesized 84 intentional designs without any human intervention, fine-tuning, sequence optimization, scoring by another algorithm or any other selection, methods that are employed in a variety of macromolecular design algorithms to wet lab workflows^4,15,16,26^. 17 of these designs were able to efficiently promote Programmed Ribosomal Frameshifting (PRF)^27,28^ and were synthesized from a variety of different prompt combinations between consensus, secondary structure, and biological function inputs. One of these synthetic RNAs, termed SS_8 efficiently changed the reading frame of the ribosome at an efficiency higher than that of many natural sequences, and has no known homology to any sequence in the universe with the computational search tools we employed.

## Results

### Constructing a semantically labeled RNA training set

A compounding factor to assessing the strength of a design model is the filtering of sequences after generation. Indeed, models generate many sequences that are then scored, ranked and handpicked with a variety of methods. For protein design models, initial designs are scored and ranked by completely different protein structure prediction algorithms^29^ such as AlphaFold^6^ or ESM^4^ followed by additional computational methods such as PLACER^30,31^. In other cases, protein design models are fine-tuned to a specific protein family for a specific design^32^. Indeed, the question arises if the design model itself has inherently learned how to design a specific sequence or if they simply act as high-volume stochastic samplers that rely on external oracles or guided fine-tuning, critical to the workflow of many design models^33^.

These bottlenecks and issues in protein design model workflows expand to full-blown obstacles in RNA design. While much work has been devoted to 3D RNA structure prediction, computational predictors still retain poor performance and are below that of human expert predictors^34^ likely due to the lack of PDB RNA structures^12^. This limits the ability of RNA design algorithms that rely on PDB structures and 3D structure predictors in their workflows^16,33^ to serve as effective designers.

Given these issues, we endeavored to train an RNA composer that required no 3D structural information and no scoring, fine-tuning, selection, ranking or hand-picking of sequences after generation.

To do this, we first focused on generating a semantically labeled training set. We turned to the RNA families database (Rfam 15^35^) as a starting point. While other RNA ML models have simply utilized the nucleotide sequences^12,19^ from Rfam and other RNA databases, we curated the specific organization of these RNAs into RNA families that group these sequences and also contain descriptions of each of these family’s functions (**Extended Figure 1A**). From these, we generated tetra-modal inputs; the RNA sequence, a consensus sequence of the RNAs within the same family, curated secondary structure annotations, and gene-ontology term(s) based on the RNA family’s biological function. The consensus sequence represents a minimized multiple sequence alignment that still contains rich evolutionary data^36^ but simplifies the training process for modality fusion. We uniformized the sequence, consensus and secondary structure for each piece of training data and then appended the gene ontology terms at the end of the vector (**Extended Figure 1B**). This database of 3,978,586 million tetramodal rich training vectors comprise semRfam with 1.21 billion multimodal tokens (**Supplementary Material**).

To properly utilize all the representations in semRfam, and because we wanted our model to reason over all these modalities simultaneously, we constructed a single and unified joint vocabulary consisting RNA nucleotides, consensus characters, secondary structure annotations and gene ontology functions.

### A model architecture that is based in the principles of RNA folding and structure

We divided the training into two phases. In the first phase, we challenged the model to learn the relationships between all the modalities and in the second phase, we then transitioned to the generative phase where the model was tasked with generating RNA sequences with none, some or all the other modalities.

In the first phase of training, where this tetra-modal training data is embedded, we also employed a 50% dropout to the consensus, secondary structure and/or gene ontology modalities. Therefore each individual training example with all four modalities is multiplied into eight with varying modalities (e.g. sequence only, sequence plus consensus, etc) which gave the model the opportunity to learn on just RNA sequences as well as every possible combination of RNA sequences with some or all modalities (**Figure 1A**). For masking we employed span masking **(Extended Figure 2)**. The architecture we employed was based on ModernBERT (**Figure 1B**). Specifically, we employed alternating global and local attention such that the model would both learn the relationship between the different modalities and long range interactions but also focus on local interactions since local interactions play a larger role in RNA structure as compared to protein structure^37^.

**Figure 1:**
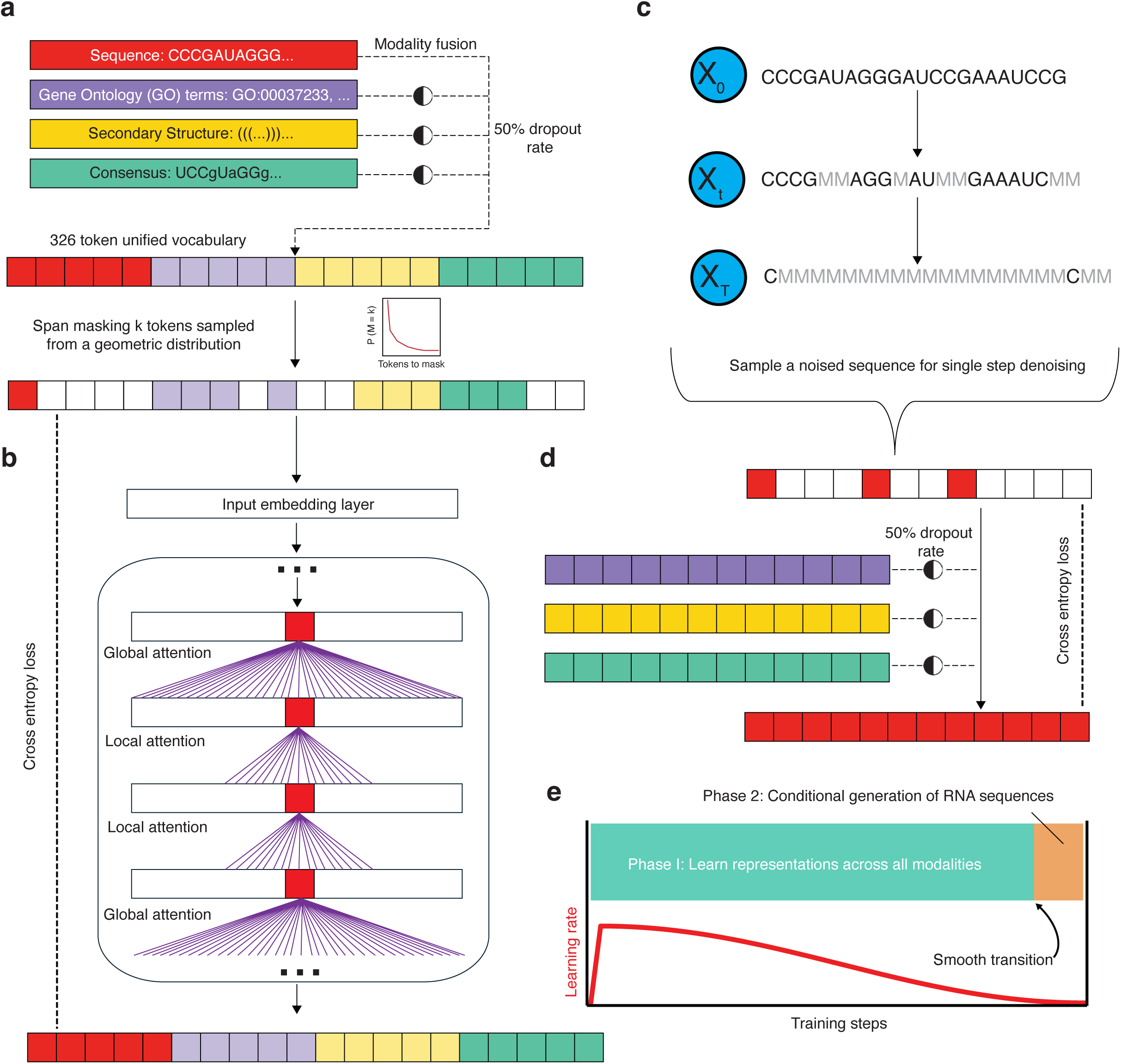
Model architecture and two stage training regime of yakRNA Design. — **a.** In the first phase of training, sequence, GO, secondary structure, and consensus modalities are fused together into a single vector with variable dropout (50%) for GO, secondary structure and consensus that are tokenized with a 326 unified vocabulary. For masking, 35% of tokens are masked using a span-based masking strategy in which spans between 2 to 8 tokens at a time are masked based on a sampling from a geometric distribution with a mean span length of 3.3 tokens. **b.** Tokenized inputs are passed through an embedding layer followed by a stack of transformer blocks that alternate between global and local attention mechanisms, enabling both long-range (intermodal and distant RNA interactions) and local range (shorter motifs) dependences to be captured. The model is trained using cross-entropy loss on all masked tokens of all modalities to reconstruct the original input. **c.** In the second stage of training, sequences are randomly noised between 0 and 95%. **d.** Now with the sequence noised randomly between 0 and 95%, the other modalities are not masked and provided with 50% dropout to regenerate the sequence with a cross-entropy loss only on the sequence. **e.** Summary of two phase training procedure with a smooth transition between learning representations between modalities to conditional generation of RNA sequence.

In the second generative phase of training, we took the RNA sequence and noised it at variable rate between 0 to 95% (**Figure 1C**). Then, the model was tasked with regenerating the sequence with none, some, or all the modalities (**Figure 1D**). This transition from the multimodal learning to generative RNA phase was smooth (**Figure 1E**) and enabled the creation of an RNA design model that can handle a variety of different modality combinations depending on the user’s specification. This yielded a model that was able to generate sequences with any combination of consensus, biological function, secondary structure, and sequence specifications desired by the user (**Figure 2A**). We split semRfam by their Rfam family designations to prevent leakage and to optimize training procedure and hyperparameters before a final production run (**Supplementary Materials**).

**Figure 2:**
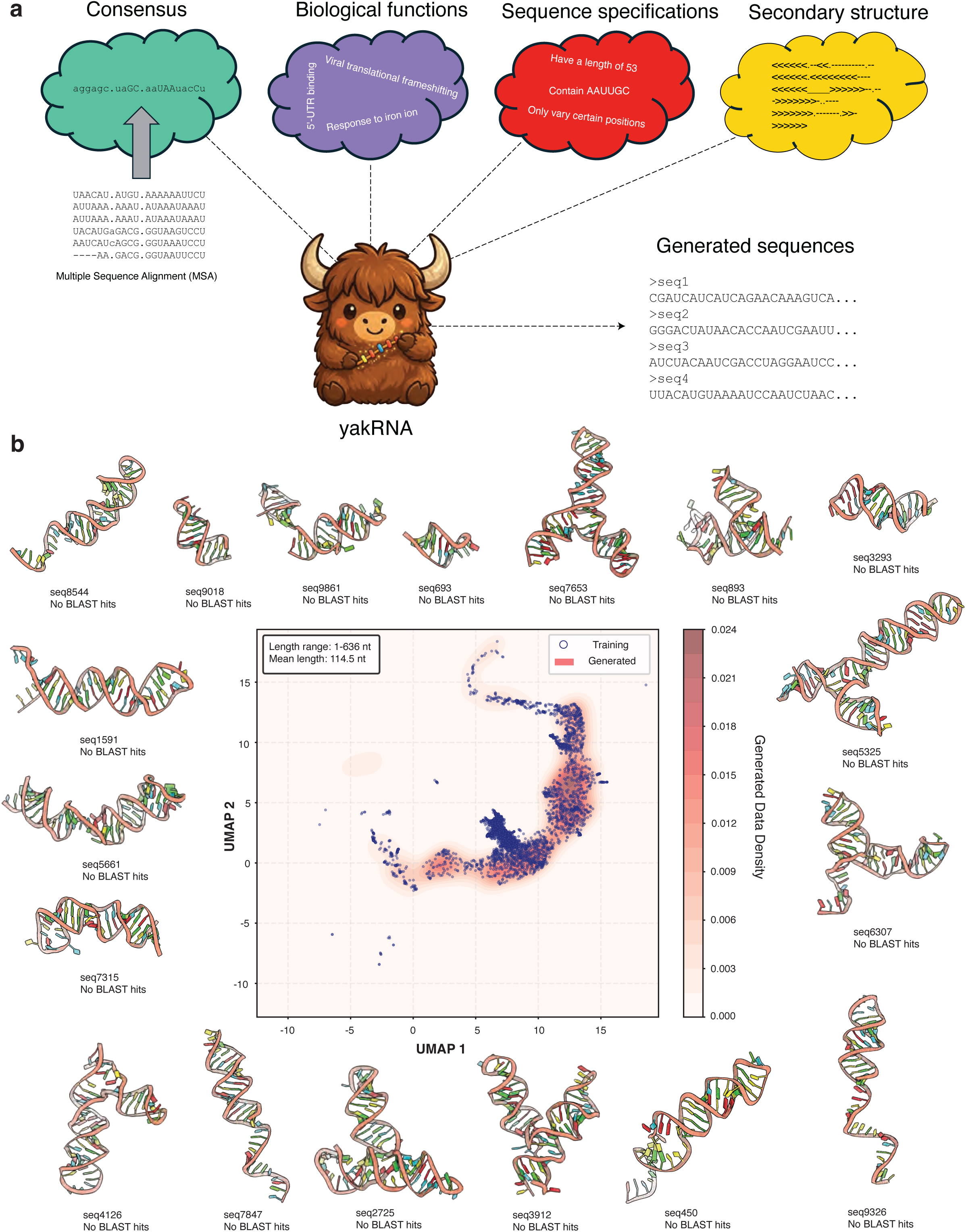
yakRNA Design can reason over sequence, biological function, secondary structure, and consensus sequence to generate plausible structured RNAs. — **a.** Schematic representation of the different prompt mixtures that yakRNA Design can take to generate sequence including consensus, biological function or GO, sequence specification, and secondary structure. **b.** yakRNA Design free generated sequences (blue dots) that were embedded along with semRfam sequences and plotted with UMAP (orange density). The max length of a sequence allowable by yakRNA design is 636 nucleotides. Sequences from this training distribution were folded with RhoFold and designs with a pLDDT greater than 70 are shown around the UMAP. These sequences were subject to BLAST but returned no hits.

After training the model, we then let the model unconditionally free generate RNAs without any prompts. RNAs generated from this model were found were able to be predicted to be structured by RhoFold^12^ and spanned the distribution of all annotated RNA families in nature (**Figure 2B**). Additionally, the majority of these RNAs contained no significant BLAST homology against nucleotide databases, suggesting that this model can generate novel RNAs and not perform simple retrieval of known sequences.

### In silico evaluation of different modalities for generation of RNAs

Given that yakRNA Design can unconditionally generate RNAs or take any combination of modalities to generate RNAs, we first wanted to assess how each individual modality shaped RNA generation and how modality combinations synergized. For this internal assessment, we examined the THF-II riboswitch family (RF02977, **Figure 3A**), which was included in semRfam during training. The THF-II riboswitch is a structured RNA riboswitch that recognizes its namesake THF ligand through a distinct RNA tertiary fold consisting of two stems connected by a central junction that regulates gene expression at the RNA level. By using a training-set family here, we can assess whether yakRNA Design has correctly internalized the relationship between modalities and their corresponding sequence space. This is a necessary prerequisite before evaluating whether the model can generalize to design functional RNAs.

**Figure 3:**
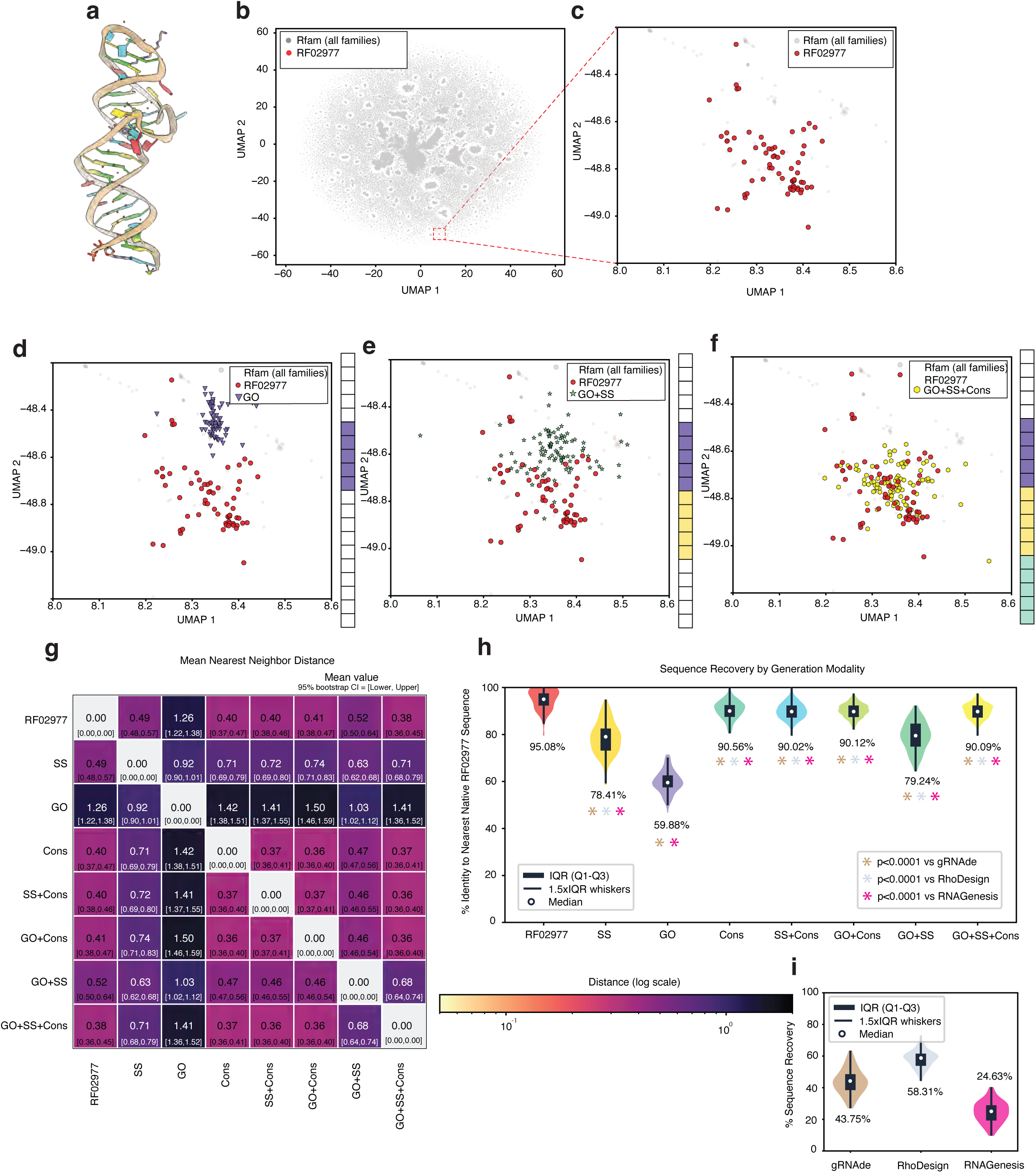
In-silico analysis of yakRNA Design’s modality synergy and comparison to other design models. — **a.** PDB model of THF-II riboswitch (PDB: 8XZP). **b.** UMAP of Rfam sequence embeddings from yakRNA Design with RF02977, THF-II riboswitch family, highlighted as red dots and other sequence embeddings as grey dots. **c.** Zoom-in of b. **d.** Embeddings from sequences generated with a gene ontology prompt shown as purple triangles in relation to RF02977 family embeddings. **e.** Embeddings from sequences with a gene ontology and secondary structure prompt shown as green stars in relation to RF02977 family embeddings. **f.** Embeddings from sequences with a gene ontology, secondary structure, and consensus sequence prompt in relation to RF02977 family embeddings. **g.** Heatmap of mean nearest neighbor distances of all modality prompt embeddings with 95% confidence interval listed below in brackets. **h.** Sequence recovery of sequences generated by each modality in relation to their most similar sequence in the RF02977 family. Sequence recovery for RF02977 generated by assessing a single sequence from the family in relation to its most similar sequence left in the family. Significance value * (p < 0.001) are presented for modalities whose sequence recovery is significantly different than those from competing models (i) assessed by an ordinary one-way ANOVA with multiple comparisons. **i.** Sequence recoveries of sequences generated by other RNA design models as compared to 8XZP’s sequence. gRNAde was prompted with both the 3D and 2D structure of 8XZP, RhoDesign was prompted with only the 3D structure of 8XZP, and RNAGenesis was only prompted with the 2D structure of 8XZP since its inverse folding of 3D structure was not accessible in the GitHub repository at the time of this assessment.

Importantly, yakRNA Design’s secondary structure modality can handle a wide array of different constraints. Despite not having 3D structural information from PDBs, the model can generate sequences with Watson-Crick, wobble, shearing and other non-canonical base-pairs because semRfam contains base pairing information of a wide variety of canonical and non-canonical interactions (**Supplementary Information**). This is important because only about ∼60% of base pairs in RNA in structured RNAs participate in canonical Watson-Crick base pairing^38^ yet many RNA algorithms only consider a very limited set of interactions^39–41^ (typically Watson-Crick plus G:U, U:G pairing).

We took all the RNAs that belonged to the THF-II riboswitch family, generated embeddings from them with yakRNA Design and then performed a UMAP^42^ to compare where the embeddings of sequences landed relative to the training set of RNA sequences (**Figure 3B-C**). Once we knew were this family landed relative to the other training sequences, we then generated sequences with yakRNA Design. We then examined the consensus, secondary structure and biological function modalities to generate sequences from scratch. While yakRNA can also take sequence specifications and perform free infilling on a partially masked sequence (**Extended Figure 3**), we did not want to provide the model with any steering sequence information to better assess how this model incorporates non-sequence modality information into design. The only sequence information provided was the length of the sequence.

First, we prompted yakRNA Design “Regulation of Gene Expression”, a phrase corresponding to GO term GO:0010468 within yakRNA Design’s vocabulary. This GO term, with regards to RNA sequences, typically annotates riboswitches. Indeed, generating sequences with just this prompt led to non-overlapping embeddings, with respect to RF02977, in the UMAP (**Figure 3D**). Next, we wanted to see if layering modalities together improved colocalization of generated sequences to the existing THF-II riboswitch sequences. We prompted the model both with the biological function and with the secondary structure corresponding to one of the THF-II riboswitches which further pushed the embeddings cluster of synthetic sequences closer to the native ones’ (**Figure 3E**). Lastly, we prompted yakRNA Design with all three of biological function, a secondary structure and the consensus sequence of the THF-II riboswitch family and generated sequences that when embedded appeared to colocalize with that of the native sequences in the UMAP (**Figure 3F**). An important computational note is that the all panels (**3A-G**) were derived from a single UMAP fit so that the relative distance comparison across the panels would be valid since UMAP distorts global distances but preserves local structure within the same UMAP run^43^.

Following these qualitative observations, we then quantitatively examined the generation of sequences from yakRNA Design with all possible modalities and compared them to each other as well as the native THF-II riboswitch sequences (**Figure 3G**) by calculating the mean nearest neighbor distance between their respective embeddings. Indeed, with regards to mean nearest neighbor distance, using all three modalities provided the smallest distance between the generated sequences’ embeddings and those of the native sequences compared to using the modalities on their own. Additionally, this analysis expectedly revealed that providing the consensus modality drove sequences’ embeddings to the native sequences’ embeddings more than the other individual modalities. Conversely, gene ontology prompting led to sequences’ embeddings that were the furthest from those of the native sequences’, perhaps owing the fact that the biological function could correspond to a variety of different RNA families.

We then performed sequence recovery analysis (**Figure 3H**). In agreement with our embedding nearest neighbor distance analysis, we found that the consensus modality provided the largest sequence recovery (90.56%) compared to the other single modalities. However, secondary structure and GO prompting still provided reasonable sequence recovery both individually (59.88% and 78.41% respectively) and together (79.24%).

Since yakRNA Design is a design model, it is difficult to use existing benchmarks used for structure and fitness comparison^44^. For the closest comparison to other models, we compared yakRNA Design to gRNAde^45^, RhoDesign^16^, and RNAGenesis^15^ which use both the 3D structure and/or secondary structure of an input to design sequences as the sole feature or one of many in the model. Both design models, trained on the PDB, and presumably contain the THF-II riboswitch in their training data, showed a mean sequence recovery of 43.75%, 58.31%, and 24.63% respectively over 100 candidate sequences per model. All of yakRNA Design’s modalities had significantly higher sequence recovery with the exception of GO only versus RhoDesign.

Following this detailed analysis on an RNA family that the model had seen, we generated 700 candidate sequences against a recently discovered RNA family, *duaA*^46^, that was annotated and deposited after this model was trained. This minimized the chance of leakage between training and test data, a persistent issue in macromolecular algorithms^47–50^. Due to the model’s robust vocabulary, the biological function annotations of these RNA sequences were interpretable by yakRNA Design. Indeed, we saw the same qualitative trend of synergy when adding modalities together (**Extended Figure 4A-E**), the consensus modality being the strongest driver of embedding distance and sequence recovery to the native Rfam family and the GO modality being the weakest (**Extended Figure 4F-G**). We were unable to benchmark RhoDesign because no 3D model exists for this brand new family but were able to use gRNAde and RNAgenesis in secondary structure mode only. gRNAde and RNAgenesis showed 27.12% and 27.34% sequence recovery respectively and all yakRNA Design modalities showed significantly higher sequence recoveries (**Extended Figure 4H**).

These results demonstrate that yakRNA Design creates sequences that faithfully recapitulates the sequence characteristics of families and that the modality inputs are correctly steering generation toward the intended RNA family. Additionally, yakRNA design’s robust vocabulary and diverse training data and regime allows it to work well on RNA families outside of its training data.

Crucially these in silico results, while demonstrating that yakRNA Design has correctly internalized the relationships and can understand RNAs outside of its training regime, cannot address whether the model can generate truly novel functional RNAs without any post-generation filtering or selection. Furthermore, sequence recovery is an imperfect metric since yakRNA Design could be designing RNAs that are diverse in sequence but equally functional to native RNAs, which would be desirable from a design standpoint.

### Zero-shot generation of functional synthetic RNAs that stimulate PRF

To assess yakRNA Design’s true generative capabilities, we sought a design challenge with stringent biochemical constraints: RNAs that must i) fold into a defined 3D shape, ii) bind to a specific molecular machine, and iii) resist active mechanical perturbation. Indeed, we challenged yakRNA Design to design synthetic RNAs that could mimic the effect of natural and complex RNA structures of Programmed Ribosomal Frameshift (PRF) signals^27,28^ whose most efficient structures consist of complex 3D pseudoknots that specifically and directly bind to the ribosome^51^ to change the ribosome’s reading frame in a programmed manner while simultaneously resisting the helicase activity of the ribosome^52^.

To prompt the model, we decided use the SARS-CoV-2 RNA frameshift stimulatory pseudoknot as our starting prompt based on the criteria we laid out above. The SARS-CoV-2 pseudoknot was shown to specifically interact with ribosomal proteins uS3, uS5, eS10, eS30 and helix h16 of the 18S rRNA to presumably promote PRF (**Figure 4A**)^51^. We prompted the model with a mix of the biological function (“viral translational frameshifting” or GO:0075523), the consensus sequence of all coronavirus pseudoknots and annotated secondary structure.

**Figure 4:**
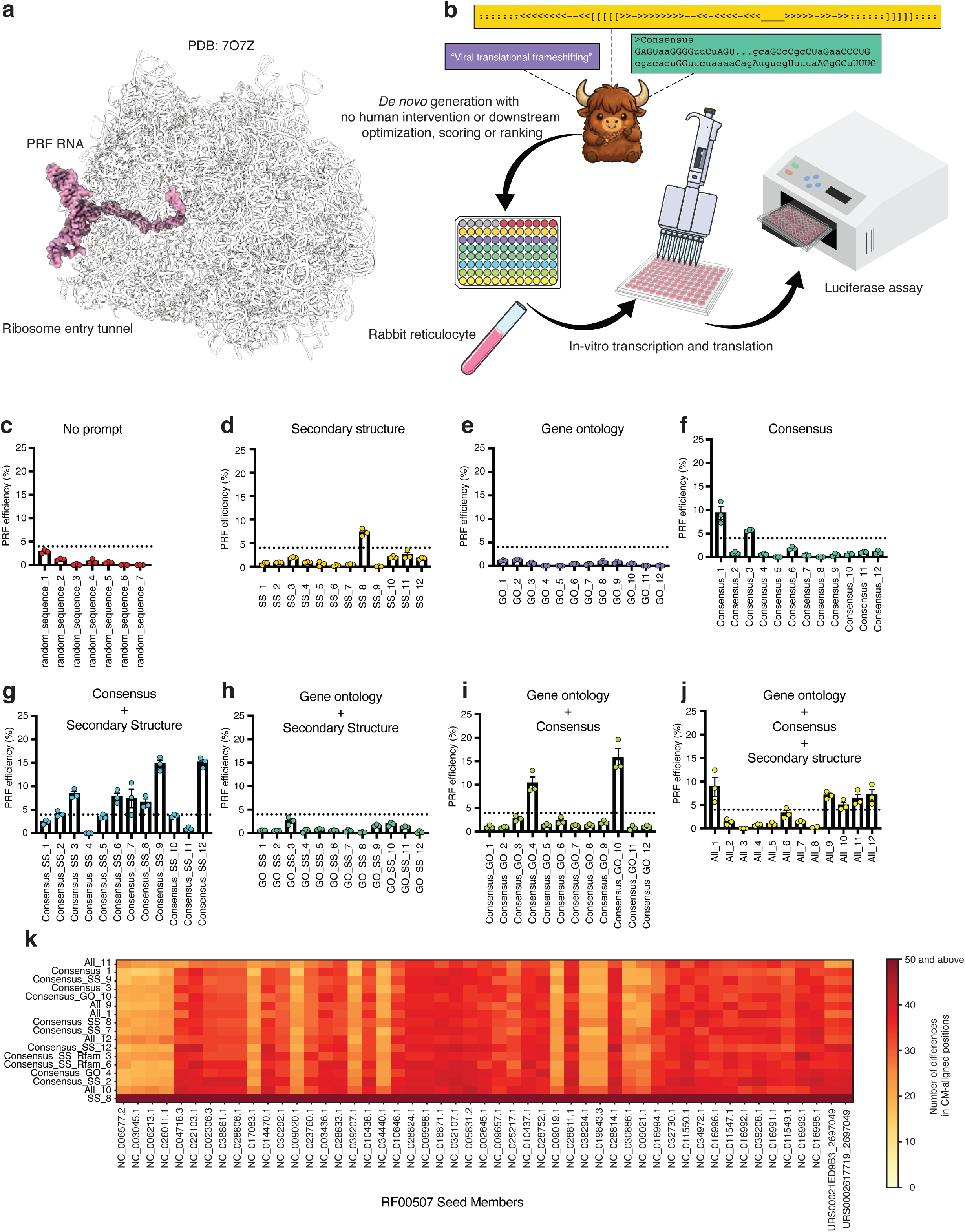
Evaluation of synthetic RNAs directly designed by yakRNA Design with no further selection. — **a.** Atomic structure of a PRF RNA pseudoknot (highlighted as pink) binding directly to the entry tunnel of the ribosome from PDB: 7O7Z. **b.** Schematic of inputs used by yakRNA Design to directly synthesize candidate structured RNAs without any additional scoring or selection criteria. Sequences were directly subject to in-vitro transcription and translation reactions that were then subject to luciferase assays. **c-j.** Programmed Ribosomal Frameshift (PRF) efficiencies of sequences generated from **c**. no prompt (only length of 78 to match other sequences), **d.** secondary structure, **e.** gene ontology, **f.** consensus sequence, **g**. consensus and secondary structure, **h.** gene ontology and secondary structure, **i.** gene ontology and consensus, and **j.** all three modalities. Dotted line represents threshold for efficient designs, which was calculated by taking the average efficiency of no prompt and multiplying it by 5. **k.** Analysis of efficient designs (y-axis) assessed by the number of differences in CM-aligned positions to seed sequences in the coronavirus RNA pseudoknot family (x-axis).

yakRNA Design directly designed a 96-well plate in which the first row contained 7 free generations, 12 secondary structure only RNAs, 12 biological function only RNAs, 12 consensus only RNAs, 12 consensus + secondary structure RNAs, 12 biological function + secondary structure RNAs, 12 biological function + consensus RNAs, and finally 12 all modality RNAs. These RNAs were directly synthesized by IDT DNA with no post-generation filtering, scoring or selection of any kind and subsequently used for an in-vitro transcription and translation assay to measure their efficacy in promoting PRF using a dual luciferase assay^53^ (**Figure 4B, Extended Figure 5A-C, Supplementary Materials**).

While some PRF RNAs stimulate efficient frameshifting at a rate as low as ∼1%^54,55^, we set our threshold at 5x the average rate of PRF efficiency for RNAs generated by yakRNA Design with no extra modality prompt, 0.8%, (**Figure 4C**) and thus an aggressively high threshold of 4% efficiency; for reference, HIV-1 has a reported PRF efficiency of as low as 2.41% in certain systems^56^ and slippery site mutations to the SARS-CoV-2 RNA, the classic negative control for PRF assays, showed a mean frameshift efficiency of 0.01% (**Supplementary Data**).

With this threshold, we found 17 bonafide synthetic RNAs that efficiently induced PRF that were among secondary structure only, consensus only, consensus+secondary structure, biological function+consensus, and biological function+secondary structure+consensus modalities with efficiency rates as high as 15.8% (**Figure 4D-J**) which is higher than reported values for IBV and SARS coronaviruses^57^, RSV^58^, HIV-1^56,59^, and other −1 PRF RNAs in nature^60–62^. As an additional control, we found that the raw firefly luciferase values were commensurate with the frameshifting efficiency (**Extended Figure 5D-K**) while the renilla luciferase values remained relative constant (**Extended Figure 5I-S**), controlling for possible reporter artefacts.

We then performed an alignment using *cmalign* (from infernal^63^) between the 17 efficient synthetic RNAs and all the seed RNAs in coronavirus pseudoknot stimulatory family (**Figure 4K**). What we found for 16 of these synthetic RNAs is that the number of differences for these 78mers or 81mers (prompts with a consensus modality where 81 nucleotides long, and all others were 78) was between 14 and 23 when compared to their respective best match within the family. For reference, single or few mutations to a typical WT PRF element can completely destroy its activity^57^. Furthermore, none of these synthetic RNAs had any significant BLAST homology, indicating that they represented unique and efficient RNAs that fit within the coronavirus pseudoknot stimulatory family just as any other newly discovered natural sequence would.

One of the synthetic RNAs, SS_8, showed no alignment to any RNAs in the seed family, BLAST homology or RNAhub homology^64^ and its embeddings did not colocalize to any natural pseudoknot’s embeddings in UMAP qualitative analysis (**Extended Figure 6A-C**). Furthermore, RhoFold was unable to predict its fold confidently (**Extended Figure 7**). Despite this, sequence analysis reveals that the RNA maintains the key structural features to form a functional pseudoknot (**Extended Figure 8A-B**) and stimulates PRF at a rates comparable or above other, natural −1 PRF elements^56,58,60^ (**Extended Figure 9A-C**).

Lastly, we evaluated yakRNA Design’s synthetic RNAs with Evo 2, a genomic foundation model trained on 9 trillion nucleotides that has demonstrated competitive zero-shot prediction of mutational effects on ncRNA fitness^65^. While Evo 2 was not trained on viral sequences, functional RNA pseudoknots are ubiquitous in cellular genomes^66^. Evo 2 delta log-likelihood scores, computed relative to a wild-type pseudoknot reference sequence, showed no significant correlation with PRF efficiency across all 91 designed and control sequences (Spearman ρ = +0.080, p = 0.453), and assigned near-identical scores to sequences spanning a wide range of PRF efficiency (**Extended Figure 10).** Thus, yakRNA Design’s ability to compose functional synthetic RNAs whose activity is not predicted by evolutionary sequence conservation underscores its generative capacity.

## Discussion

In this work, we have developed an RNA composer, yakRNA Design, that combines sequence, biological function, secondary structure, and consensus alignment to program RNAs with desired functions. Notably, our design algorithm does not use any 3D structural information in its training or a 3D structural prediction head/program to score and rank its generated sequences. Additionally, we employ no post-generation selection of sequences, which is employed by numerous other macromolecular design algorithms. yakRNA Design directly developed 84 sequences where rows of 12 sequences were synthesized with a variety of different prompts representing different modalities. Out of these 84 intentionally designed sequences, 17 of them proved to be efficient synthetic RNAs and one of these 17 showed absolutely no homology to any existing sequence, demonstrating that yakRNA Design does more than simple sequence retrieval.

Although yakRNA Design shows promising design capabilities, it has several limitations. First and foremost, although we intentionally built yakRNA Design without 3D structural information, it lacks the capability to perform inverse folding on 3D structures^67^ that other design algorithms are able to perform^16^ (though a 3D structure can be converted into a 2D representation with DSSR, that yakRNA Design can work with^68^). While the data for 3D structured RNAs in the PDB is limited, there are still examples of riboswitches and other noncoding functional RNAs that could provide models an additional modality to help a model understand how RNAs intrinsically fold and function. The issue with this approach is that the number and diversity of sequences far exceeds the number of unique structures in PDB, which creates a natural limitation on models that rely on this limited training data. Indeed, our sequence recovery metrics on an RNA family in our training data (**Figure 3H**) far exceeds the sequence recoveries of other model’s performance on RNAs in their own training data who use 3D structural data^15^. Indeed, eliminating RNA tertiary structure has shown promise in more effectively predicting RNA-drug interactions^69^. Perhaps incorporating alternative structural data such as chemical probing data^70^ could help improve predictions and design. Our approach of providing rich alternative data sources and/or vocabularies for RNA algorithms has shown promise in secondary structure prediction^23^ and 3D motifs^71^.

For the time being, it seems that leveraging RNA sequences that have been evolving for over 4 billion years^72^ and their expert, manual curation by scientists^35^ for generative models such as yakRNA Design will help drive the expansion of programmable RNAs that will benefit basic science, industry, and human health.

## Methods

The methods are available in the Supplementary Materials.

## Extended Figure Legends

**Extended Figure 1:**
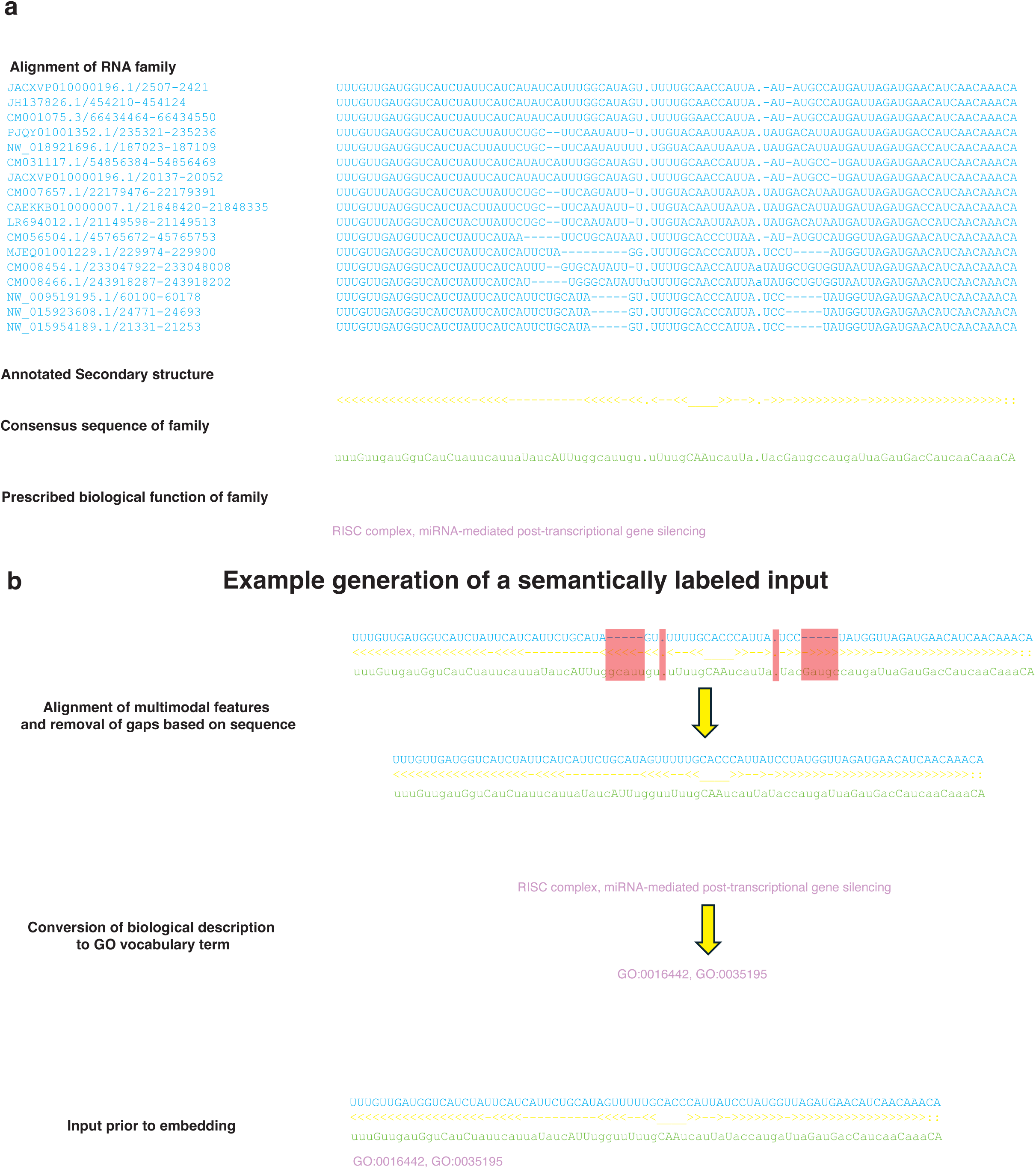
Workflow of how semRfam was constructed. — **a.** Example depiction of an aligned RNA family, annotated secondary structure, consensus sequence, and prescribed biological function. **b.** Depiction of how these separate modalities are combined to create a semantically labeled input. Red boxes represent portions that are deleted in training examples to create identical length sequence, secondary structure, and consensus sequence modalities with gene ontology terms converted to a GO accession appended to the end. The generation of these semantically labeled examples enables co-learning.

**Extended Figure 2:**
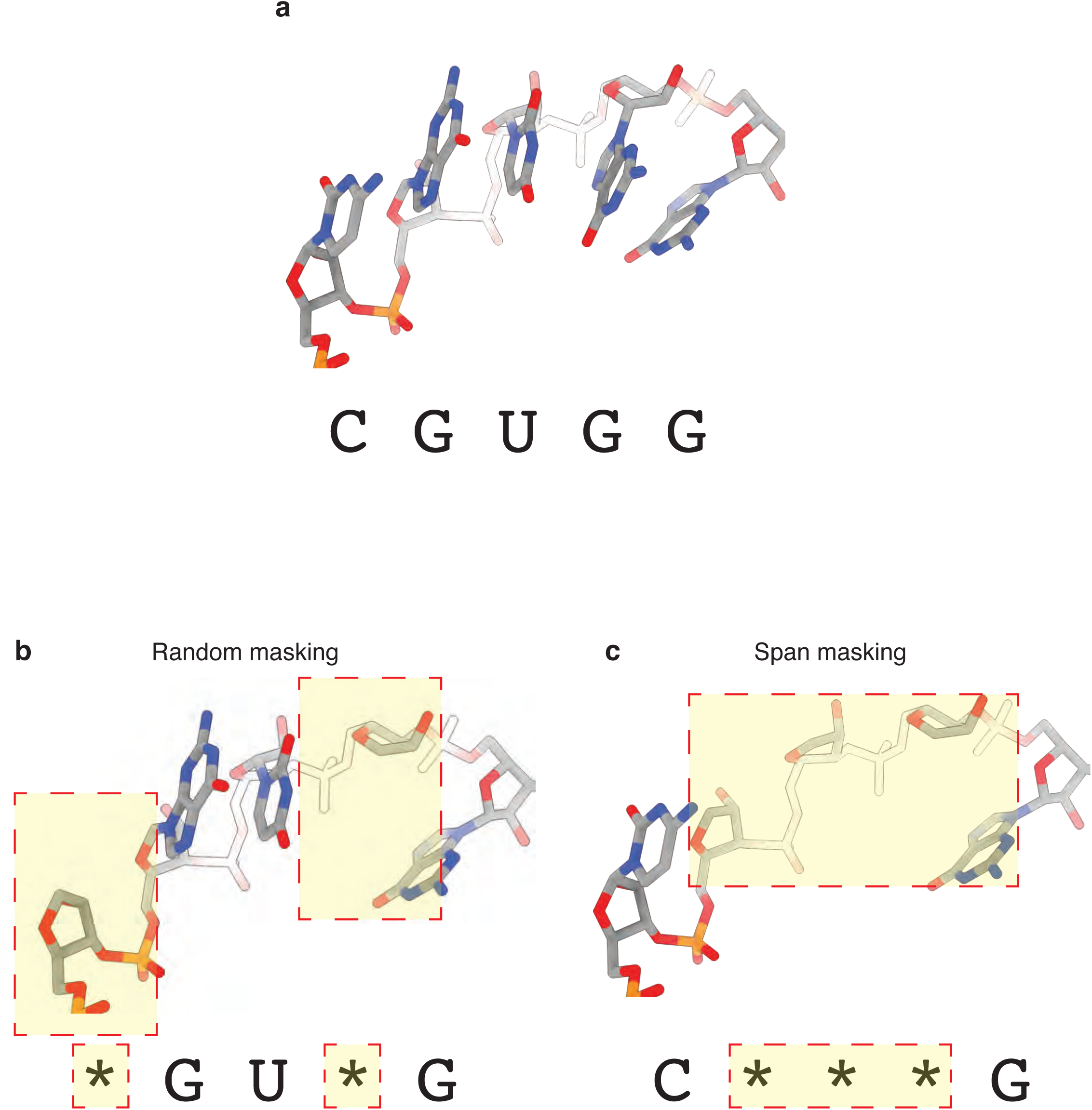
Rationale for span masking over traditional random masking. — **a.** Example of RNA in a stacking arrangement from PDB 1YHQ. **b.** Depiction of how random masking on a sequence implicitly maps onto a structured RNA. **c.** Depiction of how span masking on a sequence implicitly maps onto a structured RNA, challenging a model to learn local learn motifs.

**Extended Figure 3:**
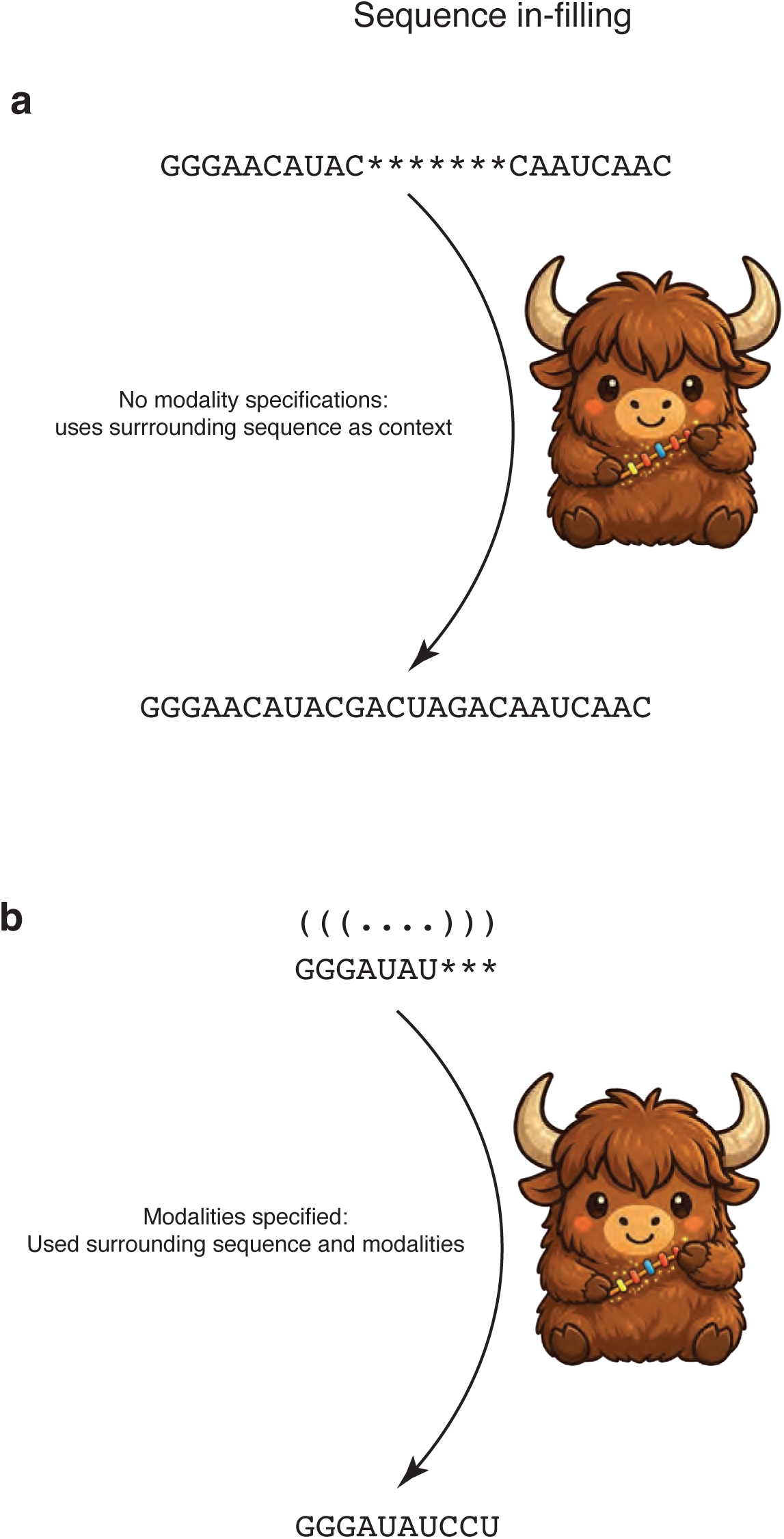
Demonstration of sequence in-filling capabilities of yakRNA Design. — **a.** Schematic demonstration of how partial sequences can be passed into yakRNA Design and have masked locations in-filled based on the surrounding sequence as context. **b.** Schematic demonstration of how both partial sequences can be passed into yakRNA Design with a combination of other modality specifications]. In the example shown, yakRNA design can both perform in-filling and faithfully follow secondary structure modality specifications.

**Extended Figure 4:**
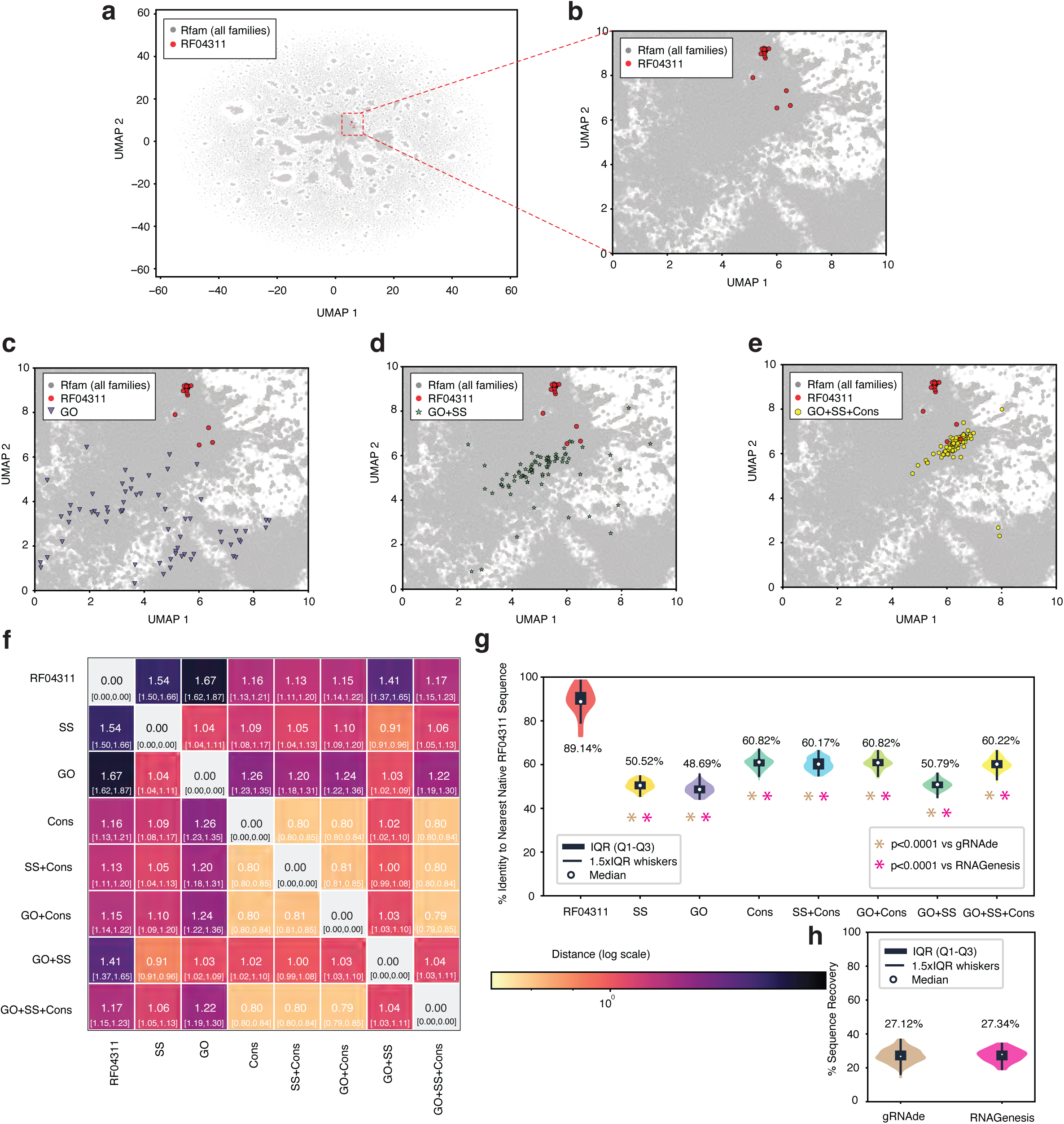
In-silico analysis of yakRNA Design’s modality synergy and comparison to other design models on a newly annotated RNA family. — **a.** UMAP of Rfam sequence embeddings from yakRNA Design with RF04311, *duaA* riboswitch family, highlighted as red dots and other sequence embeddings as grey dots. **b.** Zoom-in of b. **c.** Embeddings from sequences generated with a gene ontology prompt shown as purple triangles in relation to RF04311 family embeddings. **d.** Embeddings from sequences with a gene ontology and secondary structure prompt shown as green stars in relation to RF04311 family embeddings. **e.** Embeddings from sequences with a gene ontology, secondary structure, and consensus sequence prompt in relation to RF04311 family embeddings. **f.** Heatmap of mean nearest neighbor distances of all modality prompt embeddings with 95% confidence interval listed below in brackets. **g.** Sequence recovery of sequences generated by each modality in relation to their most similar sequence in the RF04311 family. Sequence recovery for RF04311 generated by assessing a single sequence from the family in relation to its most similar sequence left in the family. Significance value * (p < 0.001) are presented for modalities whose sequence recovery is significantly different than those from competing models (h) assessed by an ordinary one-way ANOVA with multiple comparisons. **h.** Sequence recoveries of sequences generated by other RNA design models as compared to the prompt sequence from which secondary structure was dervied. Both gRNAde and RNAGenesis were prompted with the 2D structure of a candidate *duaA* family. RNAGenesis was only prompted with the 2D structure since its inverse folding of 3D structure was not accessible in the GitHub repository at the time of this assessment.

**Extended Figure 5:**
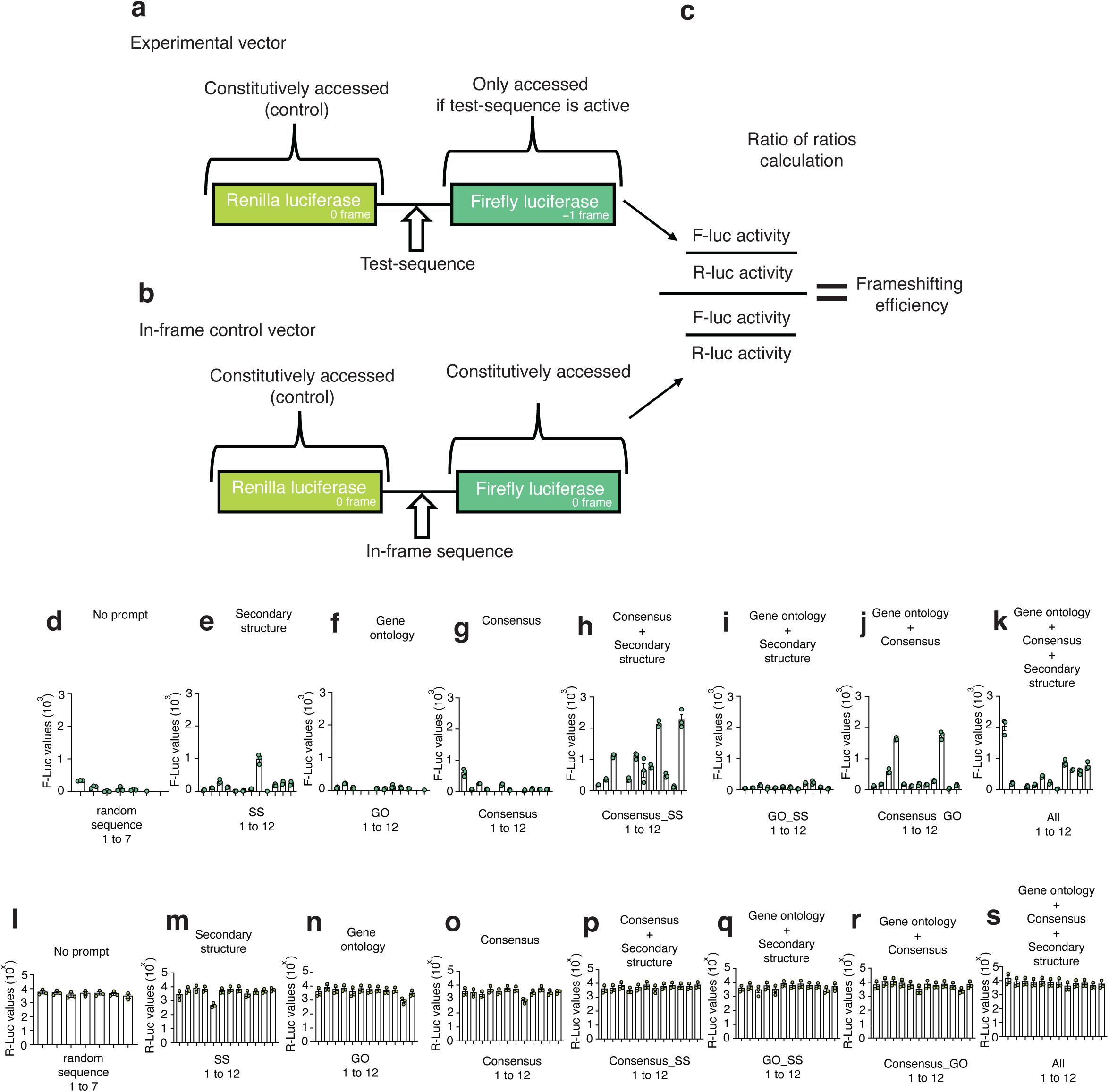
Dual luciferase assay schematic and reporter value assessment. — **a.** Schematic of the experimental vector used for assessing synthetic RNA designs from yakRNA Design. Renilla luciferase is constitutively expressed while firefly luciferase is only expressed if the synthetic RNA is active and efficiently changes the reading frame of the ribosome during translation. **b.** Schematic of the in-frame control vector which provides a baseline control of activities of both renilla and firefly luciferase. **c.** Ratio of ratios calculation to calculate the effective Programmed Ribosomal Frameshift (PRF) efficiency of a given candidate sequence. **d-k**. Raw firefly luciferase values (after subtracting water background control) from constructs with synthetic pseudoknots generated from **d.** no prompt, **e.** secondary structure only, **f.** gene ontology, **g.** consensus, **h.** consensus and secondary structure, **i.** gene ontology and secondary structure, **j**. gene ontology and secondary structure, **k.** all three modalities. **l-s**. Raw renilla luciferase values (after subtracting water background control) from constructs with synthetic pseudoknots generated from **l.** no prompt, **m.** secondary structure only, **n.** gene ontology, **o.** consensus, **p.** consensus and secondary structure, **q.** gene ontology and secondary structure, **r**. gene ontology and secondary structure, **s.** all three modalities.

**Extended Figure 6:**
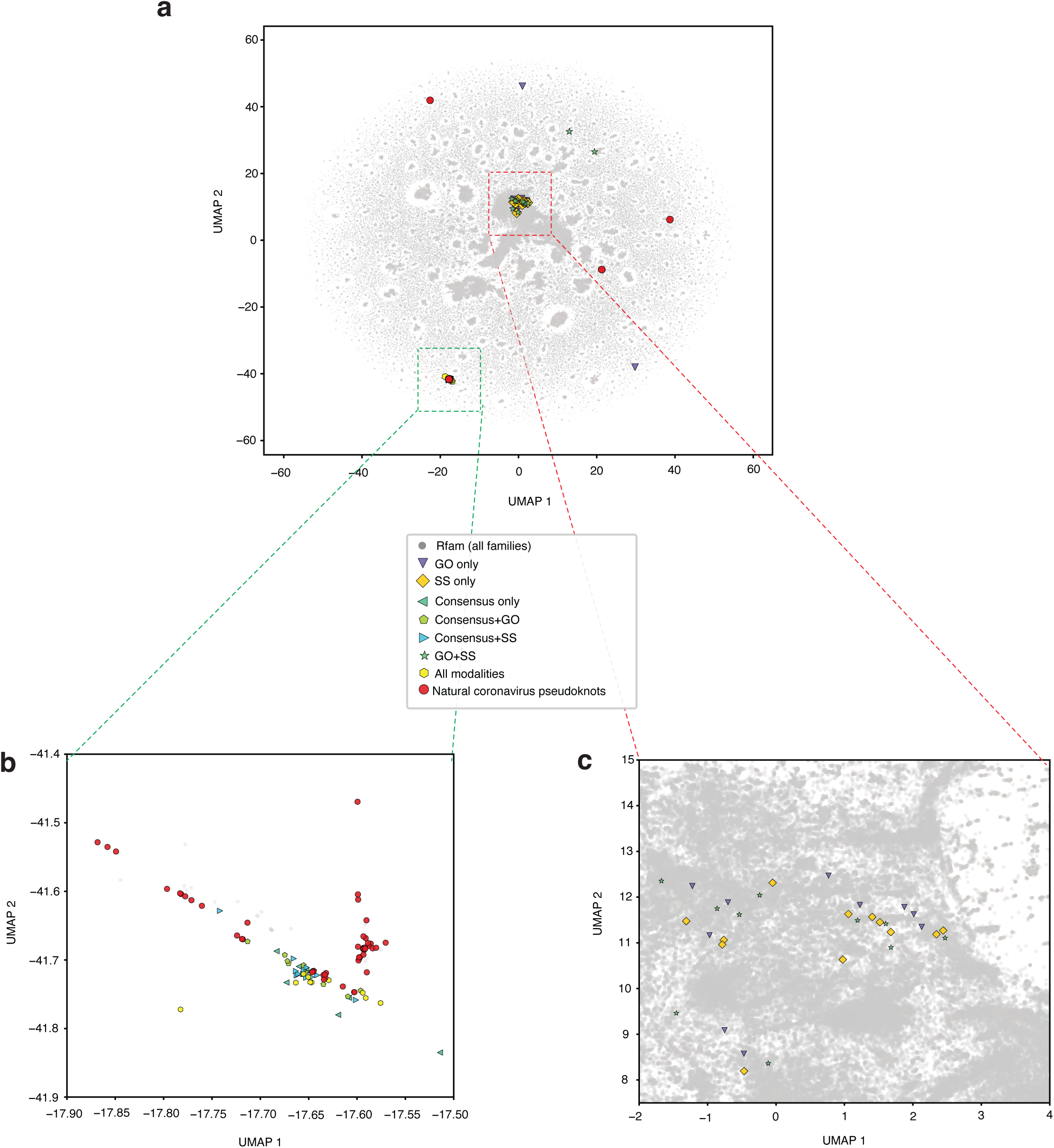
In-silico analysis of yakRNA Design’s of synthetic pseudoknots. — **a.** UMAP of natural coronavirus pseudoknots (red) and synthetic designs (see legend) as compared to all Rfam sequences (grey dots). **b.** Zoom in of majority of natural coronavirus pseudoknots (red) compared to synthetic generations. Only consensus, consensus+GO, consensus+SS, and all modalities generations are seen here. **c.** Zoom in of other synthetic generations. No natural pseudoknots are seen here but have GO, SS, and GO+SS modalities seen here.

**Extended Figure 7:**
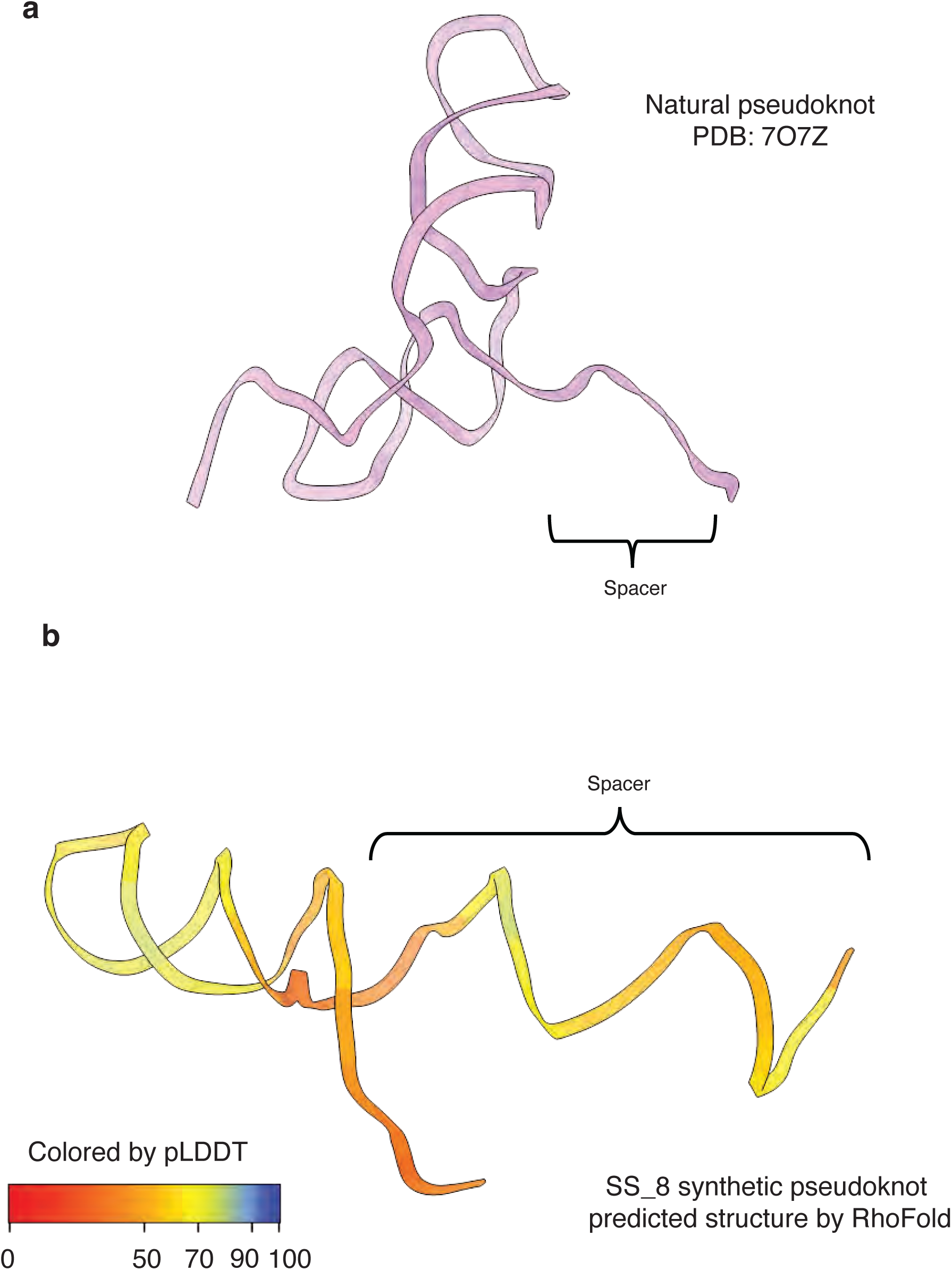
Comparison of natural pseudoknot to RhoFold prediction of SS_8. — **a.** Cartoon representation of PDB 7O7Z, specifically the pseudoknot portion. The 7 nucleotide spacer is shown in the schematic **b.** RhoFold single sequence prediction of SS_8. The overall confidence, as assessed by pLDDT, is poor across the entire structure. The putative predicted spacer region is far too long to support any sort of frameshifting since typical spacer regions must be between 5-9 nucleotides so that a downstream RNA structure may contact the ribosome while the slippery site is in the decoding center. Lastly, the predicted structure is that of a stem-loop, which is inconsistent with the specified secondary structure at the time of generation.

**Extended Figure 8:**
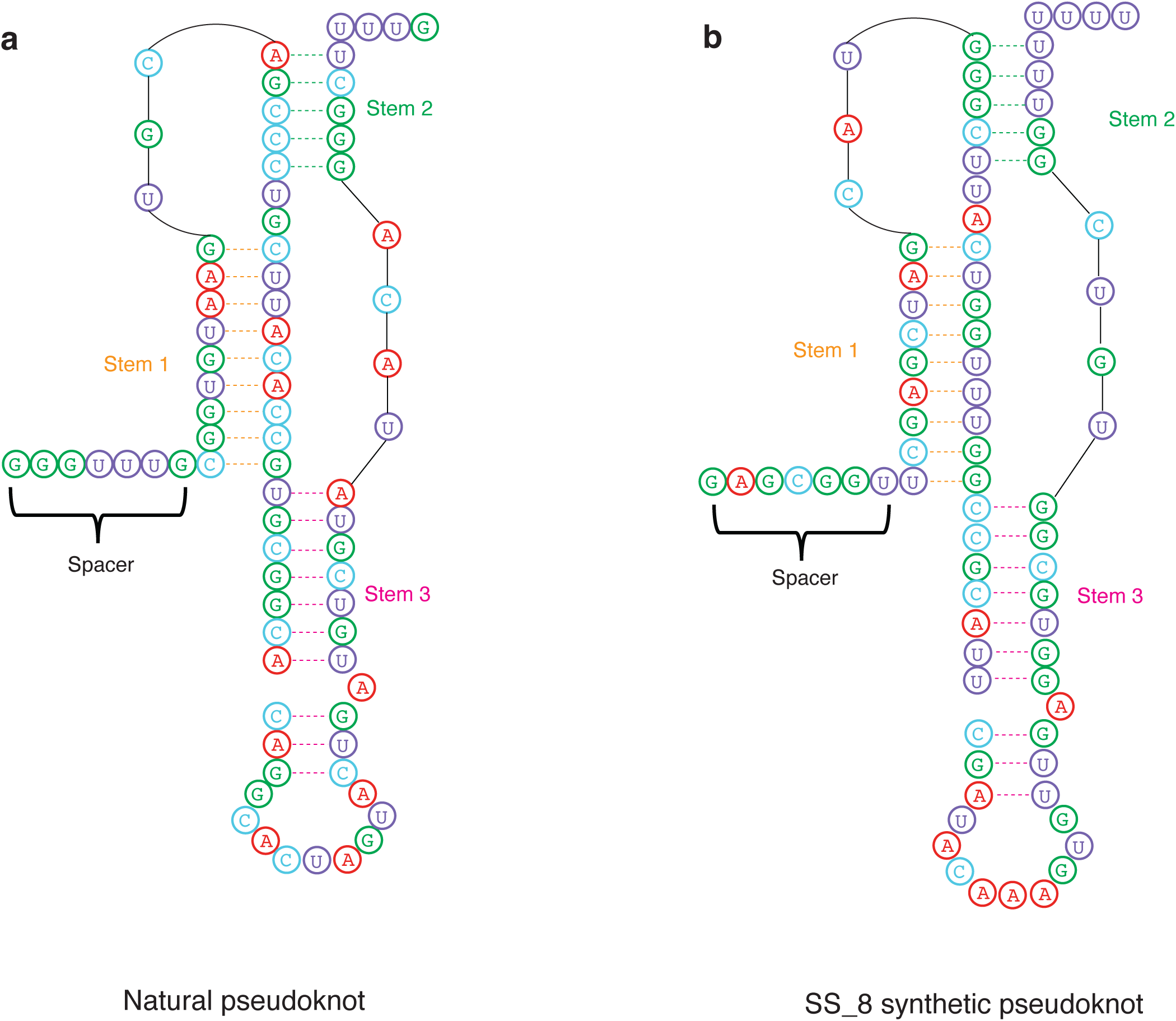
Sequence analysis of natural pseudoknot to SS_8 shows that SS_8 can adopt a pseudoknot. — **a.** Sequence analysis of the structure formed by pseudoknot in PDB 7O7Z depicting a three-stemmed pseudoknot. **b.** Sequence analysis of SS_8 pseudoknot shows that it has the requisite nucleotides at the requisite positions to fit the constraints of forming a three-stemmed pseudoknot as well, which is consistent with its high level of efficient PRF.

**Extended Figure 9:**
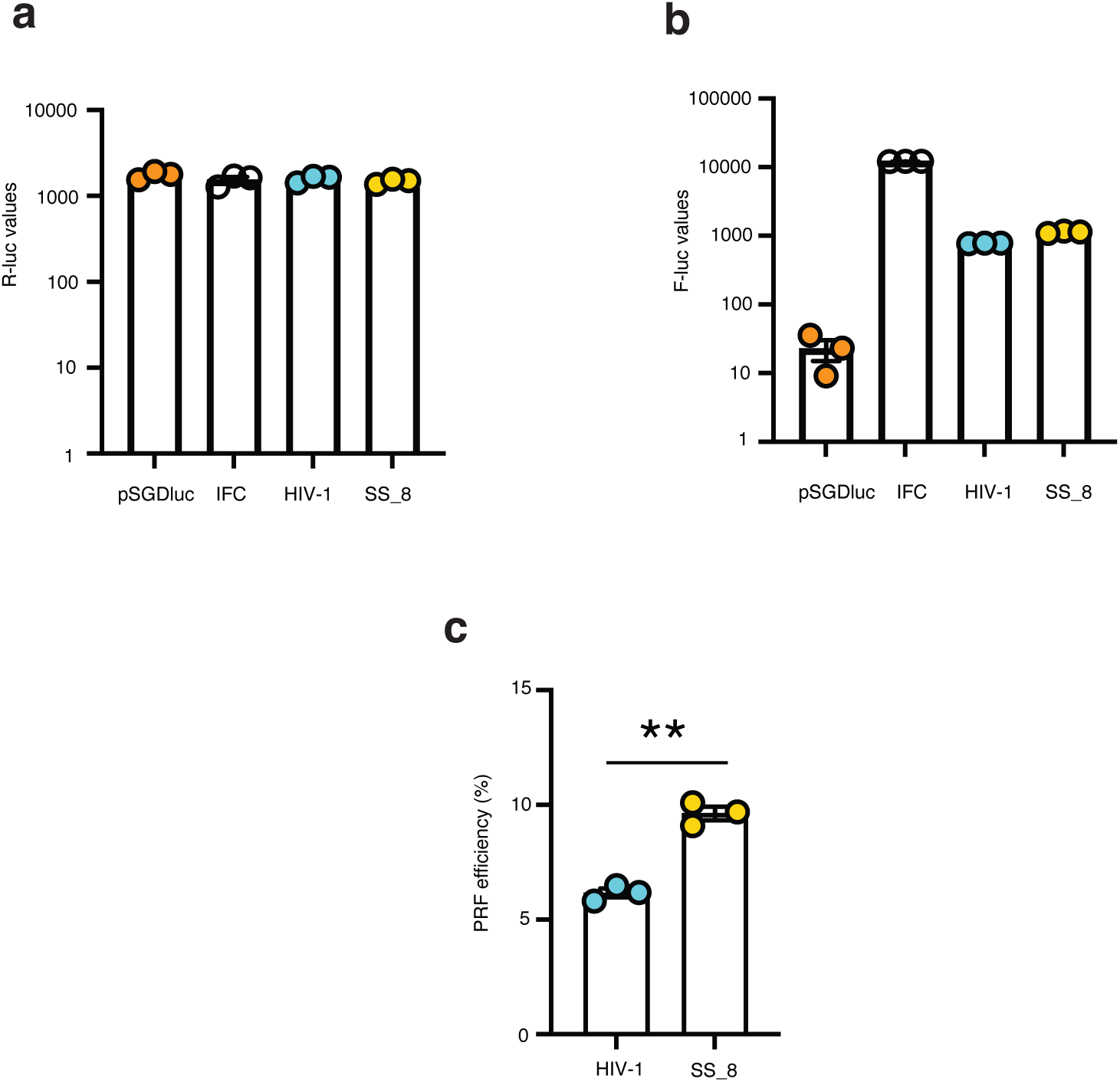
Comparison of SS_8 to HIV-1’s PRF element. — **a-c** Were conducted in a non-high throughput standard rabbit reticulocyte volume as specified by Promega. **a.** Raw renilla luciferase values from the assays conducted with the water control subtracted. **b.** Raw firefly luciferase values minus water control. **c.** Calculated PRF efficiency of HIV-1 and SS_8. ** indicates a p-value <0.01 and was conducted using a two tailed T-test.

**Extended Figure 10:**
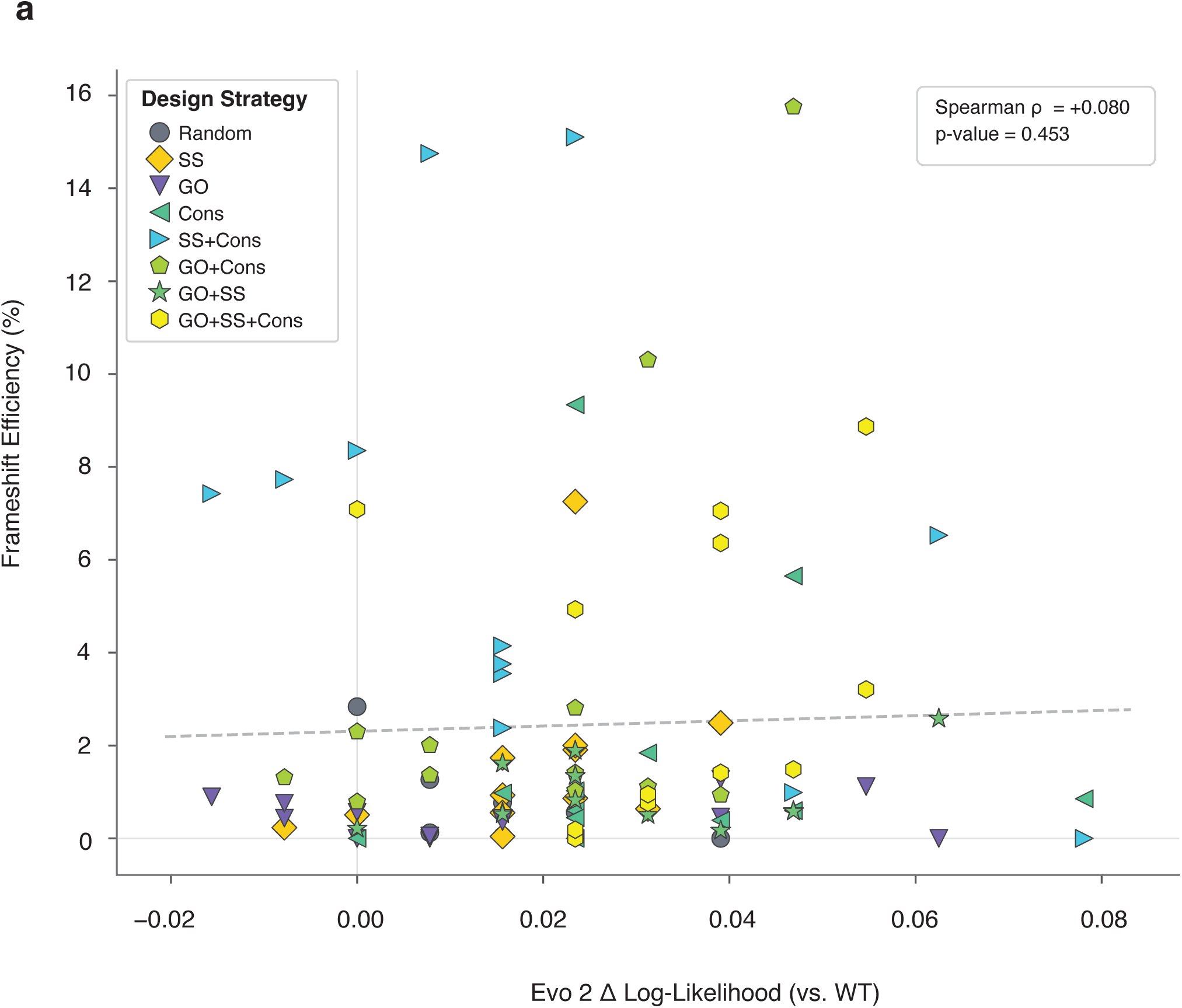
Evo 2 log-likelihood does not predicted frameshift efficiency of yakRNA Design generated sequences. — **a.** Scatter plot of Evo 2 7B delta log-likelihood (Δ log-likelihood, x-axis) versus experimentally measured programmed ribosomal frameshifting (PRF) efficiency (%, y-axis) for all 91 sequences, comprising 7 random controls and 84 sequences generated by yakRNA Design across 7 design strategy combinations. Sequences are colored and shaped by design strategy as indicated in the legend. The dashed grey line indicates the linear regression fit across all sequences. Spearman rank correlation and associated p-value are shown (inset). The near-zero, non-significant correlation (ρ = +0.080, p = 0.453) and the compressed Δ log-likelihood range (−0.02 to +0.08) across sequences spanning a wide range of PRF efficiency indicate that evolutionary sequence naturalness, as captured by Evo 2, does not predict the functional activity of yakRNA Design-generated pseudoknot sequences.

## Data availability

The model weights are available at: https://huggingface.co/MasterYster/yakRNA-Design

## Code availability

The inference code repository is available at: https://github.com/YousufAKhan/yakRNA which includes a link to an easy to use google colab notebook.

## Acknowledgements

We thank Dr. Andrew E. Firth for helpful discussions regarding the design of the frameshifting experiments and constructs. Yousuf A. Khan was funded by an NIH DP5 grant (1DP5OD039455-01), the Stanford School of Medicine as a distinguished fellow, and Dr. John L. Hennessy.

## Author contributions

Y.A.K. conceived of the project, wrote the manuscript and performed all computational work. L.N.P. and Y.A.K. planned and performed the luciferase assays. L.N.P. and Y.A.K. edited and finalized the final draft of the manuscript.

## Supplementary Materials

Supplementary tables 1-4, supplementary figures S1-S5, and methods can be found in the supplementary materials document.

