## Supplementary Materials for "yakRNA Design: A semantic multimodal RNA composer"

1. Table of Contents

### 2. Methods

#### 2.1. Training yakRNA Design

##### 2.1.1. Data Sources

Training data were derived entirely from the Rfam website and database annotations (release 15.0^1^), a comprehensive collection of non-coding RNA (ncRNA) families defined by their sequence, secondary structure, and evolutionary conservation. Two components were obtained: (1) Stockholm-format multiple sequence alignments for each Rfam family and (2) the rfam2go file mapping Rfam accessions to Gene Ontology (GO) functional annotations^1^.

Each Stockholm alignment file encodes three forms of information for a given family: the individual member sequences (with gap characters marking unaligned columns), the family-level consensus secondary structure (SS_cons annotation line), and the consensus sequence derived from the alignment's conserved positions (RF annotation line). This structure allows each training example to be simultaneously annotated with its sequence, predicted base-pairing topology, and evolutionary conservation signal — the three modalities central to this work.

GO term annotations were parsed from the rfam2go.txt mapping file, which provides cross-references between Rfam family accessions and GO identifiers across molecular function, biological process, and cellular component ontologies. These annotations introduce a fourth modality connecting sequence-structure information to functional categories.

##### 2.1.2. Preprocessing pipeline

Raw data processing proceeded in four sequential stages executed by a single pipeline script.

GO term extraction. The rfam2go file was parsed to construct a JSON dictionary mapping each Rfam accession to a set of associated GO identifiers. Lines were filtered for well-formed Rfam:RFXXXXX ... GO:XXXXXXX entries.

Stockholm parsing. For each Rfam family directory, the corresponding Stockholm file was read to extract all sequence member rows along with the shared SS_cons and RF consensus annotation lines. Each member sequence was paired with its family's consensus structure and consensus sequence, and annotated with the family's GO terms, yielding one record per member sequence with all four modalities.

Sequence cleaning and column alignment. Cleaned sequences were produced by (i) retaining only alignment columns containing alphabetic nucleotide characters, (ii) stripping gap characters, (iii) mapping non-standard IUPAC nucleotide characters to N, and (iv) uppercasing all characters to the canonical alphabet {A, U, G, C, N}. Critically, the identical column indices selected during sequence cleaning were applied to the corresponding SS_cons and consensus strings, ensuring that all three modalities remained aligned character-for-character after gap removal. This positional consistency is essential for the model to learn coherent relationships between nucleotide identity, base-pairing state, and conservation at each position.

HDF5 serialization. The cleaned, per-sequence records were stored in a compressed HDF5 file with string datasets for each modality column (sequence, ss, consensus, go_terms, rfam_id, sequence_id). GO term lists were serialized as comma-separated strings. This flat, indexed format enables efficient random-access loading during training.

A validation utility independently verified that the character distributions of cleaned sequences were confined to {A, U, G, C, N} with no residual gap or ambiguity characters.

##### 2.1.3. Unified Vocabulary Design

The central design principle of the tokenization system is a unified, offset-based global vocabulary that spans all four modalities — RNA sequence, secondary structure, consensus sequence, and GO functional annotations — within a single integer token space. Rather than maintaining separate embedding tables per modality or encoding modality identity as a learned type embedding after the fact, the vocabulary assigns each modality a contiguous, non-overlapping range of global token IDs. This ensures that any given integer unambiguously identifies both the symbol and the modality it belongs to, without requiring additional side-channel information during embedding lookup.

Special tokens occupy the lowest indices (IDs 0–8) and are shared globally across all modalities:

**Supplementary Table 1**

| Token | ID | Role |
| --- | --- | --- |
| [CLS] | 0 | Sequence-level classification token |
| [SEP] | 1 | Sequence terminator |
| [MASK] | 2 | Masked position for MLM training |
| [PAD] | 3 | Padding to batch maximum length |
| [DROPPED] | 4 | Explicitly marks a modality dropped during conditional training |
| [SEQ_START] | 5 | Delimiter opening the sequence modality block |
| [STRUCT_START] | 6 | Delimiter opening the secondary structure block |
| [CONS_START] | 7 | Delimiter opening the consensus sequence block |
| [GO_START] | 8 | Delimiter opening the GO term block |

Beginning at ID 9, four modality-specific vocabularies are placed sequentially in a fixed order. Each modality also defines modality-specific fallback tokens for out-of-vocabulary inputs:

Sequence {A, C, G, U, N, [RNA-UNK], [SEQ-GAP]} — 7 tokens. The canonical RNA nucleotide alphabet plus an N ambiguity character, a modality-specific unknown token, and a gap token.

Secondary structure {(, ), ., <, >, [, ], {, }, :, ~, -, _, ,, [STRUC-UNK], [STRUC-GAP]} — 17 tokens. The full Dot-Bracket notation alphabet covering simple stem-loops as well as pseudoknot notations used in Rfam's SS_cons annotation.

Consensus sequence {A, C, G, U, a, c, g, u, ., ~, [CONS-UNK], [CONS-GAP]} — 12 tokens. Uppercase letters denote universally conserved positions; lowercase letters denote moderately conserved positions; . and ~ denote variable or insertable positions, encoding evolutionary conservation in the token identity itself.

GO terms — Variable number of tokens, one per unique GO identifier observed in the training corpus (e.g., GO:0003735), plus [GO-UNK] and [GO-GAP] fallbacks. The total GO vocabulary size is determined empirically from the dataset during the vocabulary-building step.

The modality offset for each category is computed by accumulating the sizes of all preceding modality vocabularies, beginning at offset 9. A complete bidirectional lookup table maps every global ID to its (token_string, modality) pair and vice versa. The total vocabulary size is therefore 9 + |seq_vocab| + |ss_vocab| + |cons_vocab| + |go_vocab|, a relatively compact vocabulary given that it spans four distinct biological information domains.

##### 2.1.4. On-The-Fly Tokenization and Length Filtering

Tokenization is performed on-the-fly during data loading rather than being precomputed and stored. This design eliminates the need to store a second tokenized copy of the dataset and allows tokenization hyperparameters to be modified without reprocessing.

For each sample, GO terms are first parsed from the comma-separated HDF5 string and truncated to max_go_terms = 12 terms, consistent with the empirical distribution of GO annotations per Rfam family. The total token length (L_{total}) is then computed. Any sample exceeding max_position_embeddings (set to 1,920 tokens in cluster training configuration and 2,000 in single-GPU configurations) returns None from __getitem__ and is silently discarded by the collation function. This length filter preserves all but a small number of exceptionally large families (RF02088, RF01825) while ensuring the entire token sequence fits within the model's positional encoding capacity.

Character-level lookup is performed for the three string modalities: for each character, its local index within the modality vocabulary is found, and the global token ID is computed as modality_offset + local_index. For unrecognized characters, the modality-specific [UNK] token is substituted. GO terms are tokenized at the term level, one token per GO:XXXXXXX identifier.

The output of _tokenize_sample is a Python dictionary containing:

input_ids — torch.LongTensor of shape (L_total,) with global token IDs

attention_mask — torch.LongTensor of ones of the same length, indicating real (non-padded) positions

labels — a copy of input_ids (later modified by the masking pipeline in the collation step)

##### 2.1.5. Batch Collation and Modality Type Tracking

Variable-length samples within a batch are padded to the maximum sample length in that batch (capped at max_position_embeddings) using the [PAD] token (ID 3). Padding positions receive attention mask value 0 and label ID −100, which is excluded from the loss computation.

In parallel with padding, the collation function constructs a modality_type_ids tensor of shape (batch_size, seq_len). This tensor assigns an integer modality index (0 = sequence, 1 = secondary structure, 2 = consensus, 3 = GO terms, −1 = special/padding) to every token position by scanning each sample's token IDs for modality delimiter tokens. This auxiliary tensor is passed to the model and used during training to restrict the logit predictions at each position to only the valid token IDs of the correct modality — a vocabulary constraint (apply_vocab_constraints: true) that sets the logits of all out-of-modality tokens to −∞ before softmax, effectively preventing the model from predicting a secondary structure character at a sequence position or vice versa. This constraint substantially focuses the learning signal and removes the need for the model to expend capacity learning these trivially impossible cross-modality assignments.

##### 2.1.6. MLM masking strategy

For the masked language modeling (MLM) pre-training phase, masking is applied within the collation function after the samples have been padded and modality type IDs have been computed. The masking pipeline proceeds in two sequential stages:

Stage 1 — Modality dropout. Before any per-token masking, entire conditioning modalities may be stochastically replaced with [DROPPED] tokens. The [DROPPED] token (ID 4) is a dedicated vocabulary entry — distinct from [MASK] — that explicitly encodes "this modality was intentionally withheld." The dropout rates are configured independently per modality: in the cluster training configuration, secondary structure, consensus, and GO terms each have a 50% per-sample dropout probability. The RNA sequence modality is never dropped in the MLM phase. This stage forces the model to reconstruct masked positions in all modalities even when one or more conditioning modalities are absent, directly training the cross-modal imputation capability needed for conditional generation at inference time.

Stage 2 — Span masking. Following modality dropout, a span-based masking strategy is applied to the remaining (non-dropped, non-special) token positions. The total masking rate is set to 35% of eligible tokens. Rather than selecting positions independently at random (as in the original BERT), contiguous spans of tokens are selected, with span lengths sampled from a truncated geometric distribution:

$$l\sim Geometric\left( 0.3 \right),\quad2\leq l\leq8$$

or

$$P\left( l \right)=\frac{0.3\left( 0.7 \right)^{l-1}}{\sum_{k=2}^{8} 0.3\left( 0.7 \right)^{k-1}}$$

yielding a mean span length of approximately 3.3 tokens. Span boundaries are constrained to fall entirely within the non-special, non-dropped candidate positions, with a double-check ensuring no special tokens are inadvertently included in any span. Valid span start positions are enumerated exhaustively, then one is selected uniformly at random. Spans are selected iteratively until the total masking budget is consumed; if no valid span of the sampled length exists, individual token masking is used as a fallback.

Each selected position is then treated with the standard BERT 80/10/10 substitution rule: 80% of masked positions are replaced with [MASK], 10% are replaced with a randomly drawn non-special token from the full vocabulary, and 10% are left unchanged. The label tensor is set to the original token ID at masked positions and to the padding label (−100) at all other positions, so the cross-entropy loss is computed exclusively over the masked positions.

For validation, a stale validation option (stale_validation) seeds the random number generators for both modality dropout and span masking deterministically using the resolved training seed and a per-sample key, ensuring that the same validation samples receive the same masking pattern across all evaluation calls throughout training, enabling consistent loss tracking.

##### 2.1.7. Phase 1: Masked Langauge Modeling Loss

The MLM training objective is a modality-weighted cross-entropy loss over the masked token positions. After the collation pipeline applies modality dropout and span masking to produce a batch, the model receives the masked input token sequence and produces a logit distribution of shape (batch_size, seq_len, vocab_size) for every position. The vocabulary constraint mask is then applied, and the loss is computed as follows.

Exclusion of [DROPPED] positions. Any position where the input token is [DROPPED] — i.e., a position whose entire modality was stochastically replaced during modality dropout — is excluded from the loss entirely. This is critical: the model is not penalized for its predictions at positions it was explicitly told were withheld, and the loss signal is restricted to positions where the original information is known. Combined with the label tensor convention (non-masked positions receive label −100 and are ignored by the cross-entropy criterion), the effective loss mask is:

$${active}_{i}=\left[ y_{i}\neq-100 \right]\cap\left[ x_{i}\neq[DROPPED] \right]$$

where $y_{i}$ is the label and $x_{i}$ is the (possibly masked) input token at position (i).

Per-modality loss computation. The total loss is not computed uniformly over all active positions. Instead, the active positions are partitioned by their modality assignment (using the modality_type_ids tensor, values 0–3), and a separate cross-entropy loss is computed for each modality independently:

$$\mathcal{L}_{\mathcal{m}}=\frac{1}{\left| \mathcal{P}_{\mathcal{m}} \right|}\sum_{i\in\mathcal{P}_{\mathcal{m}}} -\log p_{\theta}\left( \hat{y_{i}}=y_{i} \mid x \right)$$

where ($P_{m}$) is the set of active positions belonging to modality ($m\in sequence, secondary\_structure, consensus, go\_terms$).

Consensus case penalty. The consensus modality receives special treatment reflecting the biological semantics of its token alphabet: uppercase letters (e.g., A, G) denote universally conserved positions, while their lowercase counterparts (e.g., a, g) denote moderately conserved positions. Predicting the correct nucleotide identity but the wrong case — e.g., predicting g when the target is G — is a strictly less severe error than a wrong nucleotide prediction entirely. To encode this, the consensus loss is computed token-wise with reduction='none', and each loss value is multiplied by a scalar penalty factor:

$$\mathcal{l}_{i}^{cons}= {CE}_{i}*\left\{ \begin{aligned} \lambda_{case} \\ 1.0 otherwise \end{aligned} \right.$$

with ($\lambda_{case}=0.5$). This is implemented via vectorized lookup tables mapping global token IDs to their lowercase and uppercase ASCII codes, enabling efficient identification of case-mismatch pairs without string operations on the GPU. The consensus modality loss is then the mean of these penalized per-token losses.

Weighted total MLM loss. The per-modality scalar losses are combined into a single training objective via a fixed weighted sum:

$$\mathcal{L}_{MLM}=\sum_{m} w_{m}\cdot\mathcal{L}_{\mathcal{m}}$$

with weights configured as:

**Supplementary Table 2**

| Modality | Weight ($w_{m})$ |
| --- | --- |
| Sequence | 0.85 |
| Secondary Structure | 0.05 |
| Consensus | 0.05 |
| GO terms | 0.05 |

The heavy upweighting of the sequence modality reflects that (i) sequence reconstruction is the primary generative objective, (ii) the sequence modality contributes the largest number of tokens per sample, and (iii) the conditioning modalities (structure, consensus, GO) carry relatively fewer tokens and serve primarily to shape the sequence representation rather than as independent generation targets.

##### 2.1.8. Phase 2: Generative phase loss

The second training phase fine-tunes the model as a generative model conditioned on the non-sequence modalities. The loss formulation is a standard cross-entropy prediction objective over all sequence modality positions.

Sequence noising. At each training step, a random timestep ($t\sim U\left( 0,T-1 \right)$) is drawn independently per sequence in the batch (T = 100). A cosine cumulative masking schedule determines the corruption probability at each timestep:

$$\bar{\alpha_{t}}=\frac{1-cos \left( \pi t/T \right)}{2}\cdot p_{target}$$

with target corruption percentage ($p_{target}=0.90$). Each sequence position is independently corrupted with probability ($\bar{\alpha_{t}})$ by replacement with the [MASK] token — an absorbing-state process with no random nucleotide substitution. Corruption is applied exclusively to the RNA sequence modality; all conditioning modalities (secondary structure, consensus, GO terms) are passed through unchanged, subject to the independent per-modality dropout applied by the ConditionalDataPreparator before noise is added.

x₀-prediction objective. The model receives as input the corrupted sequence concatenated with the (possibly partially dropped) conditioning modalities, and is trained to predict the original uncorrupted token at every sequence position — whether corrupted or not. The model learns to directly estimate the clean data from a partially noised observation at any noise level, without needing to predict intermediate noisy states. The loss is the mean cross-entropy over all sequence modality token positions:

$$\mathcal{L}_{diff} = \frac{1}{|\mathcal{P}_{seq}|}\sum_{\mathcal{i\in}\mathcal{P}_{seq}}-log p_{\theta}\left( \hat{y_{i}}=y_{i} \mid\mathbf{x}_{t} \right)$$

where $\mathcal{P}_{seq}$is the set of all actual RNA nucleotide positions (excluding special and delimiter tokens), ($y_{i}$) is the original uncorrupted token, and $\left( x_{t} \right)$ is the partially masked input at timestep (t). The vocabulary constraint mask is applied identically to the MLM phase — only sequence modality tokens are valid predictions at sequence positions — and padding positions are excluded via the standard ignore_index = -100 convention. There is no timestep-dependent reweighting of the loss. The total diffusion training objective is:

$$\mathcal{L}_{total}=w_{\text{seq}}\cdot\mathcal{L}_{\text{diff}},\quad w_{\text{seq}}=1.0$$

No modality loss weighting is applied in the diffusion phase, as the loss is restricted entirely to the sequence modality; the conditioning modalities serve only as context and are not included in the generative objective.

##### 2.1.9. Sampling Strategy

The Rfam database is highly imbalanced in family membership size — a small number of large families contain thousands of aligned sequences while the majority contribute only tens to hundreds. Uniform random sampling over all sequences would therefore cause gradient updates to be dominated by a handful of overrepresented families, biasing the model toward common RNA archetypes such as rRNA and tRNA at the expense of structurally distinct but underrepresented families. To counteract this, all batch construction was performed using a custom family-diverse sampler that enforces a strict one-sequence-per-family rule: at each gradient step, exactly batch_size unique Rfam families are drawn without replacement, and a single sequence is sampled uniformly at random from each selected family. This decouples gradient weight from family size, giving every Rfam family equal probability of contributing to any given update regardless of how many member sequences it contains.

##### 2.1.10. Model Architecture

The model is built on ModernBERT^2^ (ModernBertForMaskedLM, Hugging Face Transformers), a modern bidirectional transformer encoder that incorporates several architectural advances over the original BERT. The base-scale configuration was used for all production training:

**Supplementary Table 3**

| Hyperparameter | Value |
| --- | --- |
| Number of hidden layers | 12 |
| Hidden size | 768 |
| Number of attention heads | 12 |
| Attention head dimension | 64 (768 /12) |
| Feed-forward intermediate size | 3072 (4x hidden size) |
| Maximum context length | 1920 tokens |
| Attention dropout | 0.15 |
| Hidden dropout | 0.15 |
| Approximate total parameters | ~114M |

The token embedding table is initialized to the standard ModernBERT vocabulary and then resized to exactly match the unified multimodal vocabulary size — the total number of global token IDs spanning the nine special tokens, four modality vocabularies, and all GO term identifiers observed in the training corpus.

##### 2.1.11. Positional Encoding

ModernBERT uses Rotary Position Embeddings (RoPE) natively^3^, replacing the absolute learned positional encodings of the original BERT^4^. RoPE encodes relative position information directly into the query and key projections of each attention layer, without requiring a separate positional embedding addition to the input. This allows the model to generalize more naturally to variable-length multimodal sequences and is well-suited to the concatenated structure of the input, where the effective distance between two positions encodes both their within-modality offset and their cross-modality separation.

##### 2.1.12. Attention Mechanism

ModernBERT implements an alternating local-global attention pattern. Certain layers apply full global self-attention over the entire sequence, while alternating layers apply local windowed attention restricted to a fixed neighborhood of each token. This hybrid design reduces the quadratic attention cost for long sequences while preserving the ability to integrate information across the full multimodal context at regular intervals. Flash Attention 2 was enabled for all GPU training runs, providing memory-efficient fused attention computation in bfloat16 precision.

##### 2.1.13. Training Configuration

Two-Phase Training Schedule

Training proceeded in two sequential phases for a total of 50,000 gradient steps:

Phase	Steps	Objective

Phase 1 — MLM pre-training	45,000	Multimodal masked language modeling

Phase 2 — Diffusion fine-tuning	5,000	Discrete diffusion sequence generation

The transition between phases was smooth — the model weights trained during Phase 1 are directly inherited for Phase 2 with no re-initialization.

##### 2.1.14. Optimizer

AdamW^5^ with fused CUDA kernel (PyTorch 2.x fused=True) was used throughout both training phases:

**Supplementary Table 4**

| Parameter | Value |
| --- | --- |
| Learning rate | 1 × 10⁻⁴ |
| β₁ | 0.90 |
| β₂ | 0.98 |
| ε | 1 × 10⁻⁶ |
| Weight decay | 0.10 |

##### 2.1.15. Training/Validation/Test splitting

Splitting by Rfam Family. A central requirement for meaningful generalization evaluation of any RNA sequence model is that the validation and test sets contain sequences from families the model has never encountered during training. Because all member sequences of a given Rfam family, and consequently semRfam, share the same consensus secondary structure and GO functional annotations, a sequence-level random split would allow nearly identical structural context to appear in both training and evaluation sets, producing an optimistic and misleading estimate of generalization. To prevent this, all data partitioning was performed at the Rfam family level: the set of unique Rfam accession identifiers was treated as the atomic unit of splitting, and individual sequences were assigned to training, validation, or test sets solely according to their family membership.

Split Proportions. The unique Rfam families represented in the full processed HDF5 dataset were shuffled with a fixed random seed and partitioned into three non-overlapping subsets in a 90 / 5 / 5 ratio of Rfam families.

All sequences belonging to a given family were assigned together to whichever partition that family fell into, ensuring strict family-level non-overlap. The resulting split indices — a list of global HDF5 row indices for each partition — were serialized to 90_5_5_family_splits.json alongside the lists of family accessions assigned to each split, enabling exact reproducibility of the partition in future runs.

Production Training Mode. For the final production training run, a use_all_data_for_training mode was enabled. In this mode, the training dataset was expanded to include all sequences from all three splits — training, validation, and test families combined. Critically, the validation and test loader datasets remained defined by their original held-out family indices and were left unchanged for evaluation purposes. This design is analogous to the standard practice of retraining on the full dataset after hyperparameter selection: the model is exposed to the full breadth of Rfam structural diversity during gradient updates, while the fixed evaluation sets provide a consistent benchmark for monitoring training progress and reporting final performance. All validation metrics reported during production training therefore reflect true held-out family generalization from the original 90/5/5 split, not in-distribution performance.

Family Lookup. A pre-computed family_lookup.json file maps each Rfam accession to the list of its global HDF5 row indices. This file was constructed by a single scan of the HDF5 rfam_id field and is used both by the split assignment logic and by the family-diverse batch sampler at training time. The combination of the splits file and the family lookup fully determines the data seen by the model at every stage of training and evaluation.

##### 2.1.16. Learning Rate Schedule

A cosine decay schedule with linear warmup was applied over the full 50,000-step training run. The learning rate was warmed up linearly from 0 to the peak value over the first 1,000 steps, then followed a single cosine decay cycle to near-zero by the final step. No restarts were used.

##### 2.1.17. Hardware

All training was performed in bfloat16 mixed precision, which provides numerical stability comparable to float32 without requiring a gradient loss scaler. The model was trained on 8 × NVIDIA A100 GPUs using PyTorch Distributed Data Parallel (DDP) with a per-GPU batch size of 96, giving an effective global batch size of 768 sequences per gradient step. Gradient checkpointing was enabled throughout to reduce activation memory at the cost of an additional forward pass per layer during the backward pass. The model was compiled with torch.compile (default mode) at training startup, and a bucket-based warmup pass over the configured sequence length buckets ([704, 832, 1024, 1280, 1408, 1536, 1792, 1920] tokens) was run prior to training to pre-trigger JIT compilation for all expected input shapes and avoid compilation overhead during training steps.

##### 2.1.18. Regularization

Regularization was applied through four complementary mechanisms: (1) AdamW weight decay (0.10) applied to all non-bias, non-layernorm parameters; (2) attention and hidden dropout (0.15 each) within the transformer layers; (3) MLM modality dropout (50% independent per conditioning modality per sample), which forces the model to reconstruct masked positions under missing context; and (4) span-based masking at a 35% rate with geometrically distributed span lengths (geometric parameter p = 0.3, mean span ≈ 3.3 tokens, maximum 8 tokens), which prevents the model from exploiting positional proximity to trivially infer masked tokens.

##### 2.1.19. Sequence Length and Filtering

The maximum total token sequence length was set to 1,920 tokens, which accommodates the full four-modality concatenation — sequence + secondary structure + consensus (each equal in length) plus up to 12 GO terms plus 6 special/delimiter tokens — for all but two unusually large Rfam families (RF02088 and RF01825), which were excluded.

#### 2.2. Inference and Sequence Generation

##### 2.2.1. Input Encoding and Conditional Inputs

The user supplies any subset of four inputs — an optional partial nucleotide sequence (and/or length specification) , a target secondary structure in Dot-Bracket notation, a target consensus string, and a list of GO term identifiers — together with either an explicit target sequence length or a secondary structure (whose length implicitly determines it). These inputs are assembled by encode_sequence_input into a single token sequence using the identical layout the model was trained on:

[CLS] [SEQ_START] ⟨sequence⟩ [STRUCT_START] ⟨SS⟩ [CONS_START] ⟨cons⟩ [GO_START] ⟨GO⟩ [SEP]

Every position in this layout is simultaneously assigned a modality index that is later written into a modality_type_ids tensor with the same semantics used in training (0 = sequence, 1 = secondary structure, 2 = consensus, 3 = GO terms, −1 = special/delimiter).

Sequence block. For de novo generation, the sequence block is filled with copies of the [MASK] token (global ID 2), each labelled with modality index 0. For infilling, the caller may pass a partial sequence string in which the wildcard character * denotes positions to be generated: fixed positions are tokenized to their corresponding global sequence IDs (A, C, G, U, or N via the modality offset), while wildcard positions are replaced by [MASK]. Unrecognized characters are mapped to [RNA-UNK]. This unifies de novo generation and structure-/sequence-constrained infilling under a single code path.

Conditioning blocks. The three conditioning modalities are encoded character- or term-wise against their modality-specific vocabularies, with fallback to the modality-specific [*-UNK] tokens for characters not in the vocabulary. For each conditioning modality that the user leaves unspecified, the corresponding block is filled with the [DROPPED] token copies for ss and consensus (to preserve positional alignment with the sequence block), and a single [DROPPED] token for go_terms. This exactly reproduces the stochastic modality-dropout behavior the model encountered in training, so that no conditioning pattern presented at inference is out-of-distribution relative to training.

##### 2.2.2. Constraint modes for Secondary Structure at inference

**Supplementary Table 5**

| Mode | Allowed pairs | What it permits |
| --- | --- | --- |
| strict | A:U, U:A, G:C, C:G | Watson-Crick only — no wobble |
| canonical | + G:U, U:G | Watson-Crick + G:U wobble (default) |
| canonical+sheared | + G:A, A:G | + sheared G:A (common in loops/tetraloops) (default for constrained generator) |
| canonical+common | + A:C, C:A | + A:C mismatches (seen in ribozyme active sites) |
| permissive | + U:C, C:U | + U:C mismatches — all frequently observed pairs |
| unconstrained | all 16 (A/U/G/C × A/U/G/C) | Model's learned distribution is fully in control at every position |

#### 2.3. Sequence design

##### 2.3.1. THF-II Riboswitch

Synthetic THF-II riboswitch (RF02977) sequences were generated using yakRNA Design’s inference infrastructure with the following combination of terms

- Biological function: GO:0010468
- Secondary structure: <<<<-------<<<<<<<______>>>>>>>->>>>:::
- Consensus sequence: CCGUUCAACUCGUucCcuCcuuGAagGgaACUACGGGAG

##### 2.3.2. *duaA* ncRNA

Synthetic *duaA* (RF04311) sequences were generated using yakRNA Design’s inference infrastructure with the following combination of terms

- Biological function: GO:0010468
- Secondary structure: (((((,....,,,<<<<-<<<<<<<---..<<.<<<<<<<____..>>>>>.>>-->>--..>>>.>>>>-..>>>>...,<<<____._____>>.>,)))))::::
- Consensus sequence: acgCCC....cUGcCGGgagcgcgcCaa..gG.guGcCgGGAcu..CcGgC.accaCcgc..gcg.cgcuG..CCGg...acGGUCuu.ccGAACC.gAGGcguCAUG

##### 2.3.3. RNA pseudoknots

Synthetic pseudoknot sequences were generated using yakRNA Design’s inference infrastructure with the following combination of terms

- Biological function: GO:0075523
- Secondary structure A (from Rfam’s annotation, for all designs numerically designated 1-6): :::::::<<<<<<<<--<<[[[[[>>->>>>>>>>--<<-<<<<-<<<____>>>>>->>->>::::::]]]]]::::
  - Longer secondary structure when used with consensus: :::::::<<<<<<<<--...<<[[[[[>>->>>>>>>>--<<-<<<<-<<<____>>>>>->>->>::::::]]]]]::::
- Secondary Structure B (directly derived from analysis of structure in PDB 7O7Z): :::::::<<<<<<<<<---[[[[[-->>>>>>>>><<<<<<<<<<_________>>>->>>>>>>::::]]]]]::::
  - Longer secondary structure when used with consensus: :::::::<<<<<<<<<-:::--[[[[[-->>>>>>>>><<<<<<<<<<_________>>>->>>>>>>::::]]]]]::::
- Consensus sequence: GAGUaaGGGGuuCuAGU...gcaGCcCgcCUaGaaCCCUGcgacacuGGuucuaaaaCagAugucgUuuuaAGgGCuUUUG

#### 2.4. Benchmarking calculations

##### 2.4.1. Qualitative UMAP

For each experiment, generated sequences from every conditioning modality combination (native family reference, SS, GO, Cons, SS+Cons, GO+Cons, GO+SS, GO+SS+Cons) were embedded and saved as .npy files (one per condition).

To ensure that Rfam and generated points live in a common 2D manifold, a single joint UMAP was fit on the vertically stacked matrix of Rfam + generated embeddings. UMAP was run with n_components=2, n_neighbors=15, min_dist=0.1, random_state=42, and the library default number of optimization epochs, using either umap-learn on CPU (low_memory=True) or the GPU-accelerated cuML UMAP (with output_type='numpy').

##### 2.4.2. Mean nearest neighbor calculations

For every pair of embedding groups A and B (e.g. two generation conditions, or a generation condition versus the native Rfam family), we computed an asymmetric mean nearest-neighbor distance in the 768-dimensional [CLS]-token embedding space. Pairwise Euclidean distances were obtained with scipy.spatial.distance.cdist; for each point in A we retained only the distance to its single closest neighbor in B, and the reported statistic is the mean of these per-point minima. To keep the calculation tractable for large groups, each group was randomly sub-sampled to at most 1,000 embeddings without replacement before distances were computed. All distances were evaluated directly in the original embedding space; the 2D UMAP (above) was used only for visualization and not for these calculations.

Bootstrap confidence intervals. 95% confidence intervals around each mean nearest-neighbor distance were obtained by a non-parametric percentile bootstrap. On each of 500 iterations, groups A and B were independently resampled with replacement to their original sizes, the same sub-sampling cap and nearest-neighbor averaging were reapplied, and the resulting value was stored as one bootstrap replicate. The reported CI is given by the 2.5th and 97.5th percentiles of this bootstrap distribution. Self-comparisons along the diagonal of the condition × condition matrix were fixed to zero and excluded from bootstrapping, because independently resampled copies of the same group no longer share point identity and would produce a spurious non-zero interval.

##### 2.4.3. Sequence recovery

For each generation condition (native family reference and each modality combination: SS, GO, Cons, SS+Cons, GO+Cons, GO+SS, GO+SS+Cons), generated RNA sequences were read from their respective FASTA files, upper-cased, converted to RNA alphabet (T→U), and stripped of alignment gap characters (., -). Each query sequence was then compared against all native sequences of the target family (e.g. RF02977 for the THF-II analysis) using a global Needleman–Wunsch alignment implemented with Biopython's Bio.Align.PairwiseAligner (mode = global; match = +1, mismatch = 0, gap open = –1, gap extension = –0.1). Percent identity was defined as the number of exact matched positions within the aligned region divided by the alignment length, and each generated sequence was scored by its best (maximum) percent identity across all native references. When the query set itself was the native family, the query was excluded from its own reference list so that the reported value reflects intra-family diversity rather than trivial self-identity.

##### 2.4.4. gRNAde^6^

THF-II Riboswitch^7^ evaluation: Inverse folding was performed on the experimental structure 8XZP using gRNAde's 3D mode, with an additional secondary structure constraint derived from DSSR^8^ analysis of the PDB structure. 1000 candidate sequences were generated across a temperature range of 0.1–1.0, from which 100 were randomly sampled for analysis. Native sequence recovery was calculated as the fraction of positions matching the 8XZP native sequence.

*duaA* ncRNA^9^ evaluation: To evaluate gRNAde's performance conditioned solely on secondary structure, a representative sequence from the RF04311 Rfam family Stockholm alignment was selected as the design target. The corresponding secondary structure was extracted from the Stockholm consensus annotation and converted to standard dot-bracket notation. 1000 candidate sequences were generated across a temperature range of 0.1–1.0, from which 100 were randomly sampled. Native sequence recovery was calculated against the selected RF04311 representative sequence.

##### 2.4.5. RhoDesign^10^

THF-II Riboswitch evaluation: Inverse folding was performed using RhoDesign. The experimental structure 8XZP was provided as input to the inference_without2d.py script, which conditions sequence generation solely on the 3D backbone coordinates without requiring a secondary structure input. 100 independent samples were generated at a temperature of 1.0 using a bash script, with each run producing a designed sequence and a native sequence recovery rate computed directly by the model. Results were saved to a CSV file for downstream analysis.

##### 2.4.6. RNAGenesis^11^

Sequences were generated using RNAGenesis with inference-time tree search guidance directed toward a target secondary structure. The target dot-bracket structure was used to score candidate sequences at each denoising step via a combined metric of base-pair distance and structural similarity. Generation used 100 DDIM denoising steps with top-p sampling (p = 0.95, η = 1.0), a tree search active size of 1 and branch size of 4, producing 100 candidate sequences per target. Sequence recovery was assessed by comparison to the original, representative sequence used.

##### 2.4.7. Evo 2^12^

All 91 sequences were scored using Evo 2. Inference was performed using the evo2 Python package with PyTorch 2.5.1 (CUDA 12.1) on a single NVIDIA GeForce RTX 4090 GPU (24 GB VRAM) in bfloat16 precision. Prior to scoring, RNA sequences were converted to DNA notation (U→T) and cloning-associated dinucleotide tails present on all designed sequences were removed. Per-token mean log-likelihood was computed for each sequence using the standard autoregressive formulation, score(S) = (1/N) Σ log P(s_{t+1} | s_1, ..., s_t), via a single forward pass through the model with log-softmax applied over the output logits and appropriate autoregressive position shifting. A delta log-likelihood (Δ) was computed for each sequence relative to a wild-type pseudoknot reference as Δ(S) = score(S) − score(S_WT), such that Δ = 0 by definition for the reference. Spearman rank correlation between Evo 2 Δ log-likelihood scores and experimentally measured PRF efficiency was computed using SciPy.

#### 2.5.  Luciferase assays

##### 2.5.1. Vector

The primary vector used for all luciferase assays in this article is pSGDluc v3, an updated version of pSGDluc^13^ that has additional splice sites knocked down. pSGDLuc v3 is an upgrade over traditional luciferase vectors because it incorporates self cleaving inteins between the renilla and firefly luciferases allowing for an “apples to apple” comparison regardless of insert (**Supplementary Figure S5**)

##### 2.5.2. Cloning

Sequences were ordered and cloned by IDT DNA using their minigene service. Plates were received already suspend at 100 ng/μL per well.

##### 2.5.3. Transcription and translation reactions

TnT Quick Coupled Transcription/Translation System (L1170) for Eukaryotic Cell-Free Protein expression was used (Promega) for all luciferase assays. The protocol was modified for a high-throughput format with the following: 400 μL of TnT quick master mix was mixed with 10 μL of 1 mM methionine. Then, 4.1 μL of this mix was added to a 96 well plate and 1 μL of 100 ng/ μL of plasmid was added to each well. This mix was incubated at 30° C for 60-90 minutes before being used for downstream analysis.

##### 2.5.4. Luciferase readings and calculations

For each well, 20 μL of LARII and then later 20 μL of S+G mix was added to each well for luciferase assay readings. Reading were taken on a Molecular Devices Spectramax M2e.

### 3. Supplementary Figures

##### 3.1.1. S1: Training set details

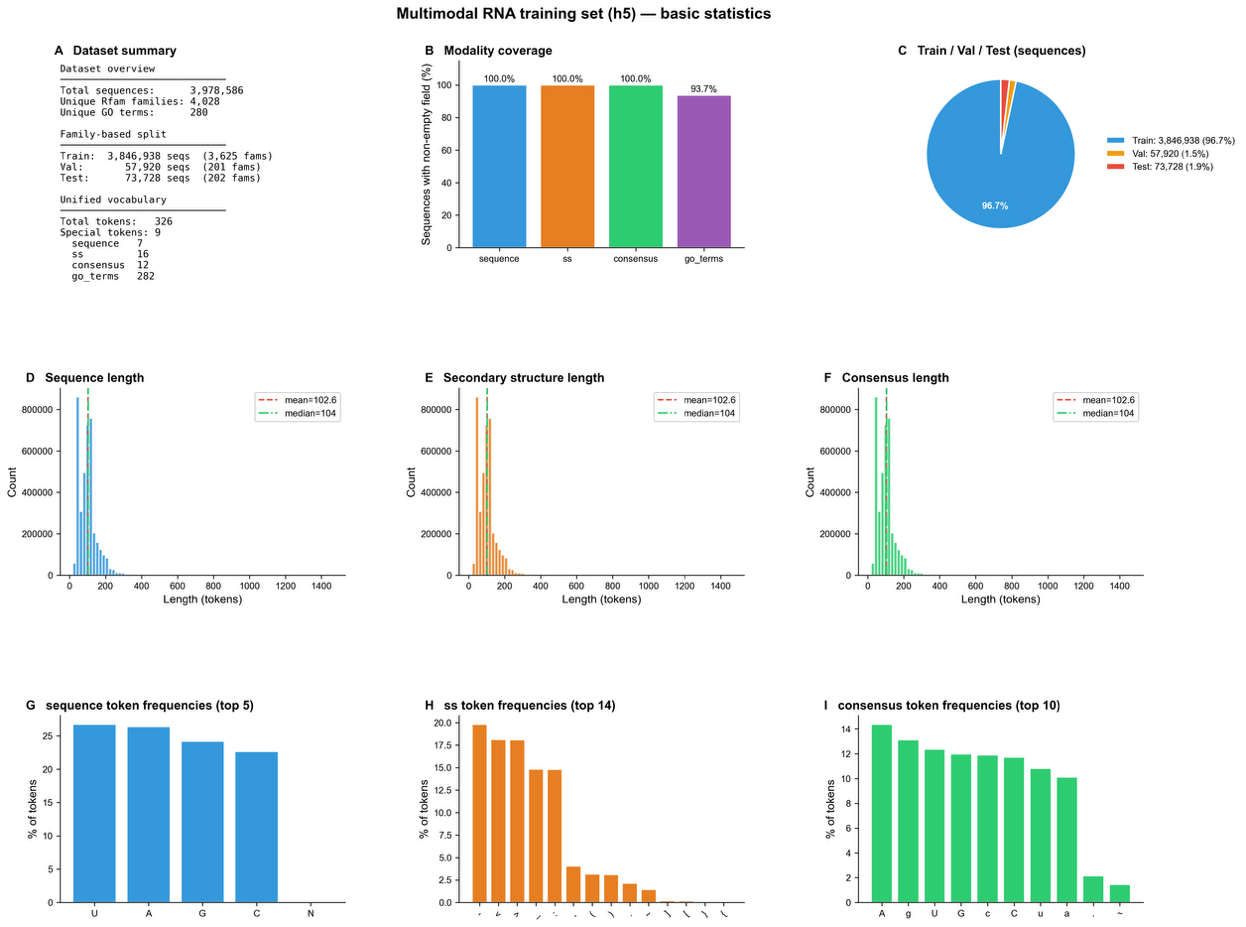

Supplementary Figure S1: Training data overview — **a.** Dataset summary of semRfam and family based splitting for training/validation/test (90/5/5). **b.** Modality coverage of each semantically constructed training example. **c.** Training/validation/test splits based on the number of sequences in the 90/5/5 family based splitting. **d.** Sequence length statistics. **e.** Secondary structure length statistics. **f.** Consensus length statistics. **g.** Sequence token frequencies. **h.** Secondary structure token frequencies. **i.** Consensus token frequencies.

##### 3.1.2. S2: Rfam family and GO term distribution

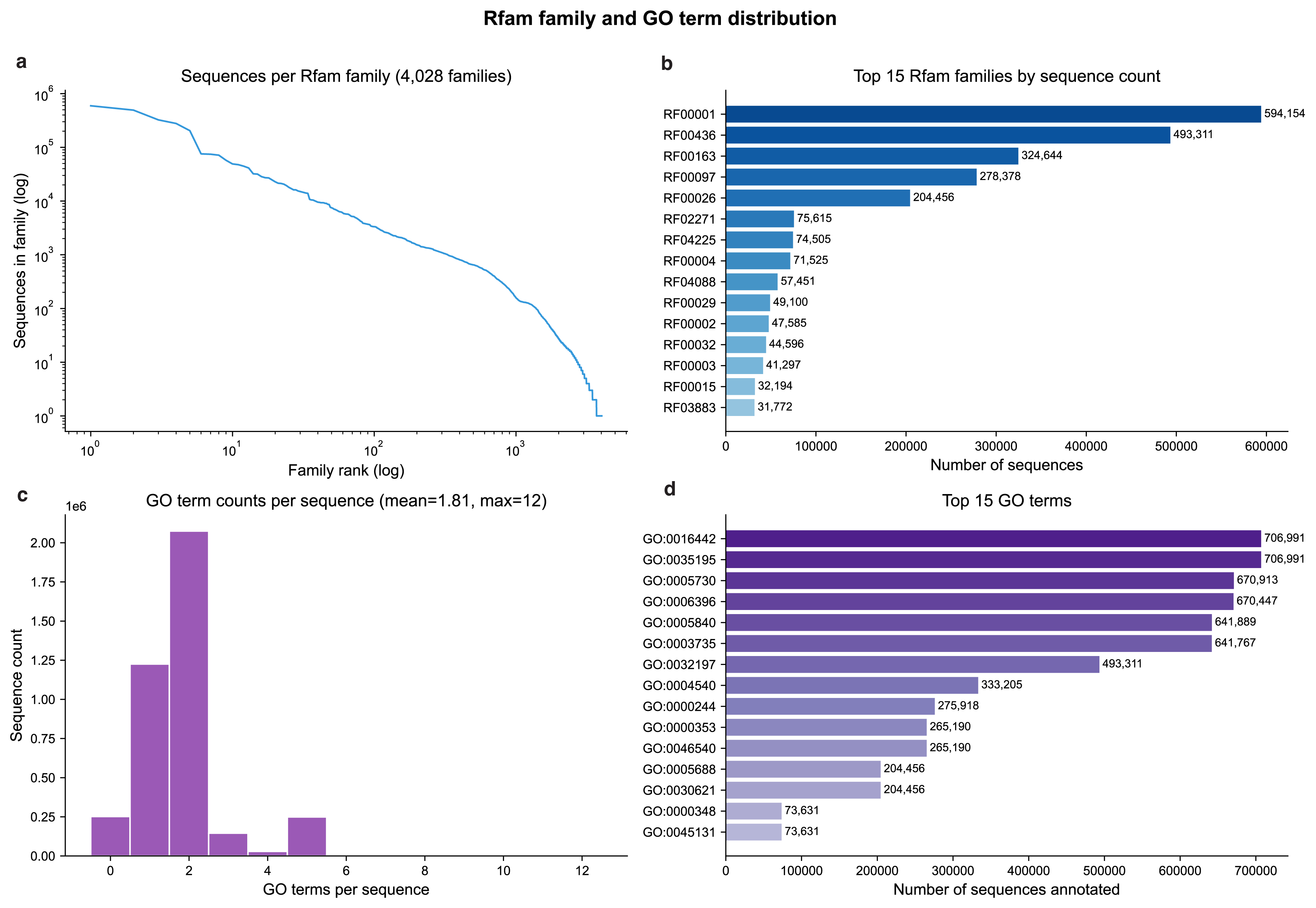

Supplementary Figure S2: Rfam family and GO term distribution — **a.** Sequences per Rfam family. **b.** Rfam families by sequence count. **c.** GO term counts per training example. **d.** Top 15 most GO terms in the semantically labeled data.

##### 3.1.3. S3: Base-pairing information

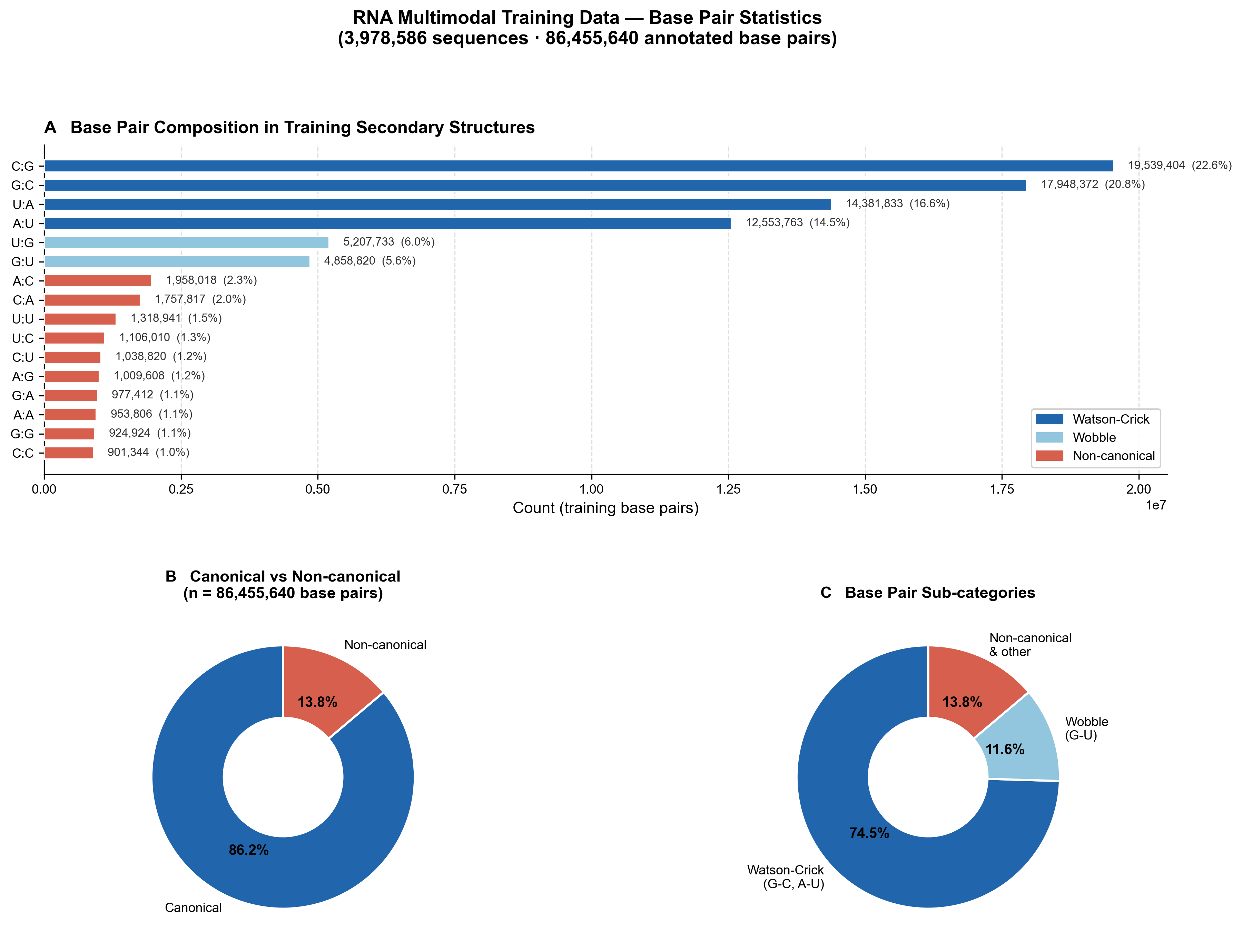

Supplementary Figure S3: Base pairing information — **a.** Base pair composition in annotated secondary structure within training data. **b.** Canonical versus non-canonical base pairs. **c.** Further sub-categorization of base pairs.

##### 3.1.4. S4: pSGDluc v3 vector schematic

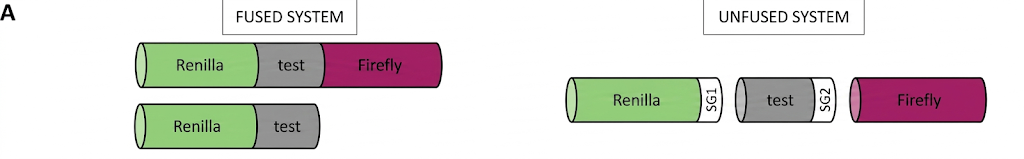

Supplementary Figure S4: Schematic of pSGDluc — **a.** Original dual luciferase assay construct. **b.** pSGDLuc v3 with self cleaving inteins.

##### 3.1.5. S5: Luciferase assay controls

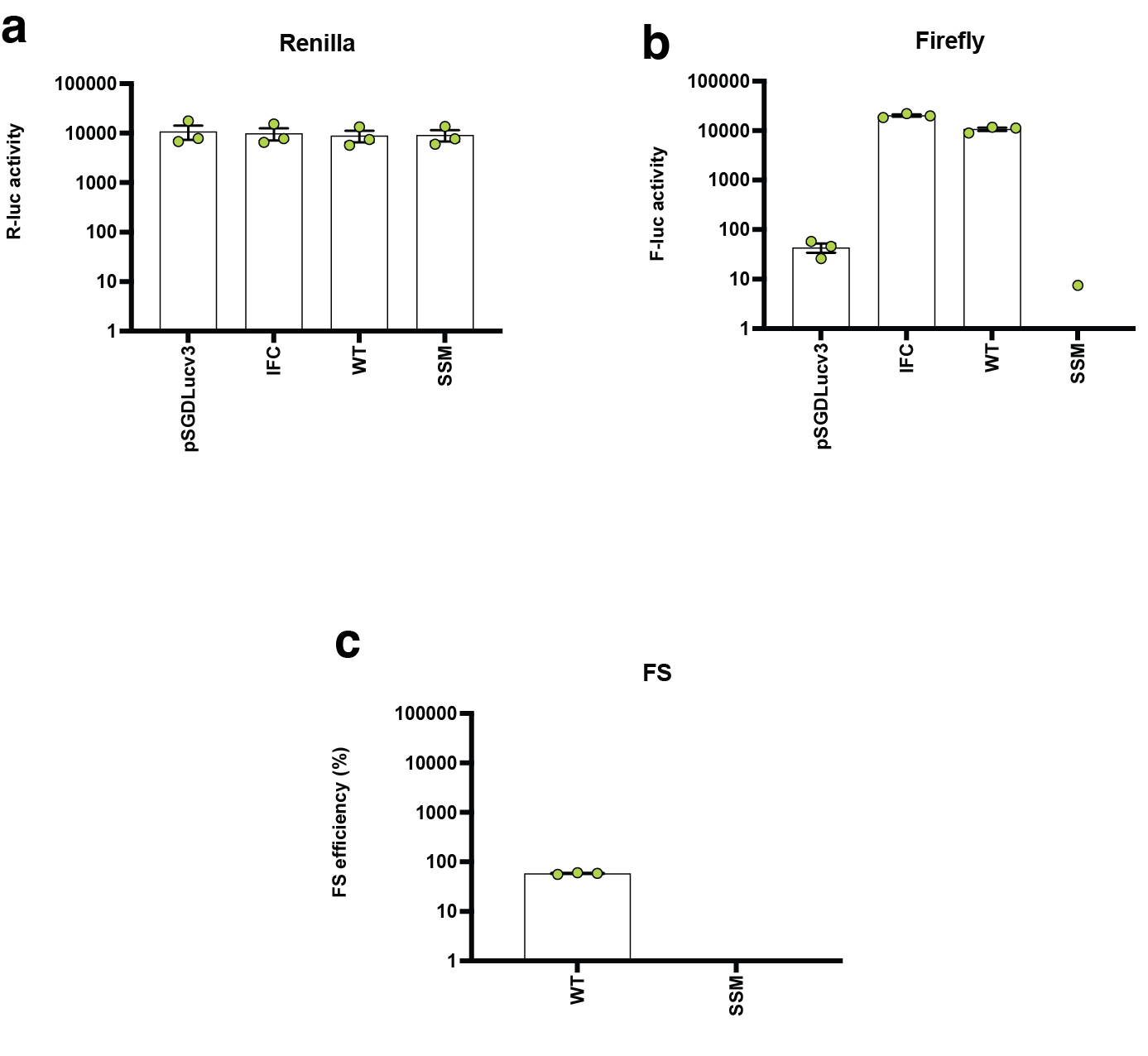

Supplementary Figure S5: High throughput luciferase assay controls — **a.** Raw luciferase values with water subtracted. **b.** Raw firefly luciferase values with water subtracted. **c.** Frameshifting efficiency for SARS-CoV2 pseudoknot, one of the most efficient coronaviruses, and its commensurate slippery site mutant.

### 4. Safety statement

The maximum length of sequence generatable is 636 nucleotides and yakRNA Design is trained to design structured RNAs, not coding ones. Additionally, given that the smallest RNA virus is ~2 kb^14^, this completely prevents this model from being used for purposes other than the generation of structured RNAs for biotechnological purposes.
